# Systematic spatio-temporal mapping reveals divergent cell death pathways in three mouse models of hereditary retinal degeneration

**DOI:** 10.1101/554733

**Authors:** M.J. Power, L.E. Rogerson, T. Schubert, P. Berens, T. Euler, F. Paquet-Durand

## Abstract

Calcium (Ca^2+^) dysregulation has been linked to neuronal cell death, including in hereditary retinal degeneration. Ca^2+^ dysregulation is thought to cause rod and cone photoreceptor cell death. Spatial and temporal heterogeneities in retinal disease models have hampered validation of this hypothesis.

We examined the role of Ca^2+^ in photoreceptor degeneration, assessing the activation pattern of Ca^2+^-dependent calpain proteases, generating spatio-temporal maps of the entire retina in the *cpfl1* mouse model for primary cone degeneration, and in the *rd1* and *rd10* models for primary rod degeneration. We used Gaussian process models to distinguish the temporal sequences of degenerative molecular processes from other variability sources.

In the *rd1* and *rd10* models, spatio-temporal pattern of increased calpain activity matched the progression of primary rod degeneration. High calpain activity coincided with activation of the calpain-2 isoform but not with calpain-1, suggesting differential roles for both calpain isoforms. Primary rod loss was linked to upregulation of apoptosis-inducing factor (AIF), although only a minute fraction of cells showed activity of the apoptotic marker caspase-3. After primary rod degeneration concluded, caspase-3 activation appeared in cones, suggesting apoptosis as the dominant mechanism for secondary cone loss. Gaussian process models highlighted calpain activity as a key event during primary rod photoreceptor cell death.

Our data suggests a causal link between Ca^2+^ dysregulation and primary, non-apoptotic degeneration of photoreceptors and a role for apoptosis in secondary degeneration of cones, highlighting the importance of the spatial and temporal location of key molecular events, which may guide the evaluation of new therapies.

## 1. Introduction

The primary light-detecting cells in the retina are rod and cone photoreceptors (rods and cones). Generally, cones are responsible for day time vision, while rods are responsible for vision under dim light conditions. Rods and/or cones degenerate and die in a heterogeneous group of genetic diseases called hereditary retinal degeneration (RD). The most common disease within this group is Retinitis Pigmentosa (RP), which first causes the loss of rods, followed by secondary cone loss. Approximately one in 3,500 individuals is affected by RP (Bertelsen et al., 2014), which presents in humans as a progressive loss of night vision, gradual constriction of the visual field (“tunnel vision”), and eventually complete blindness (Hartong et al., 2006). A related retinal dystrophy is achromatopsia, affecting about one in 30,000 individuals (Remmer et al., 2015), in which typically only the cones are lost.

Autosomal recessive forms of RP in humans are caused, for instance, by mutations in the rod-specific phosphodiesterase 6B (*PDE6B*) (Danciger et al., 1995; Hamel, 2006; McLaughlin et al., 1993). Conversely, mutations in the cone-specific *PDE6C* are a hallmark of achromatopsia (Sundaram et al., 2014). The function of PDE6 is to hydrolyse cyclic guanosine monophosphate (cGMP) after light detection, initiating the photoreceptor signalling cascade. Loss of PDE6 activity in either rods or cones results in higher-than-normal levels of intracellular cGMP, leading to constitutive opening of cyclic-nucleotide-gated (CNG) channels (Kohl et al., 2012; Thiadens et al., 2009). This in turn leads to a cation influx carried mostly by Na^+^ and Ca^2+^ (Arango-Gonzalez et al., 2014; Michalakis et al., 2018)

Strict regulation of the intracellular Ca^2+^ concentration ([Ca^2+^]) is essential for the function and survival of neurons and therefore tightly controlled by several mechanisms across cellular compartments (as reviewed in Krizaj & Copenhagen, 2002). Dysregulation of [Ca^2+^] has been theorized to initiate photoreceptor cell death, but the underlying mechanisms are still debated. For instance, high [Ca^2+^] has been proposed to trigger apoptosis (Orrenius et al., 2003), however, apoptotic caspases do not appear to be activated in most animal models for RD (Arango-Gonzalez et al., 2014; Doonan et al., 2005). Of note in this context is the mitochondrial membrane protein apoptosis-inducing factor (AIF). Despite its name, AIF is associated with non-apoptotic, necrosis-like forms of cell death (Shang et al., 2014), as its function does not require caspase activity (Bano & Prehn, 2018) and has previously been linked to cell death in photoreceptors (Sanges et al., 2006).

Strong indication for Ca^2+^ involvement in photoreceptor cell death comes from the finding of stalwartly increased calpain activity as a common denominator in many RD models (Arango-Gonzalez et al., 2014). Calpains comprise a family of Ca^2+^-activated cysteine proteases with 15 isoforms identified to date (Nemova, Lysenko, & Kantserova, 2010). Some are ubiquitously expressed throughout the body (*e.g.* calpain-1, −2), others are tissue-specific (*e.g.* calpain-3, −8) (Suzuki et al., 2004). While calpain-1 is considered to be neuroprotective, calpain-2 may be involved in neurodegeneration (Baudry & Bi, 2016). The two isoform’s Ca^2+^ affinities may indicate their presumed functions: calpain-1 activates at around 5 to 50 µM, which is much less than the ∼1 mM needed for calpain-2 activation (Khorchid & Ikura, 2002). Hence, excessive calpain-2 activation as a consequence of [Ca^2+^] dysregulation has long been linked to cell death mechanisms, but its exact role is still controversial (reviewed in Liu, Van Vleet, & Schnellmann, 2004; Rizzuto et al., 2003; Baudry & Bi, 2016).

To elucidate the degenerative progression, we performed spatio-temporal mapping in the RD and wt retina to assess the levels of calpain activity, as well as the expression of several other enzymes related to cell death, including calpain-1, calpain-2, AIF and caspase-3. To study primary cell death of both cone and rod photoreceptors, we employed the cone-photoreceptor-function loss (*cpfl1*) mouse model for achromatopsia (Chang et al., 2009) and the retinal degeneration mouse models *rd1* (Keeler, 1966) and *rd10* (Chang et al., 2002), both considered to be models for RP. In *rd1* and *rd10*, in addition to primary rod loss, we also studied secondary degeneration of cones. In contrast to the classical statistical approaches employed in previous studies, we applied a novel probabilistic modelling approach, to infer the likely sequence of degenerative events in the different mutant models, and to assign these sequences a numerical probability. Our data suggests that during the initial stages of degeneration in RP, calpain-2 is strongly activated, while calpain-1 activation appears later. This points at the execution of non-apoptotic cell death mechanisms during primary rod degeneration. However, a delayed activation of caspase-3 in *rd1* cones suggests that secondary cone degeneration is instead caused by “classical” caspase-driven apoptosis.

## 2. Material and Methods

### Animals

To study primary cone and rod degeneration, *cpfl1* (C57BL/6J background) and *rd1* (C57BL/6J x C3H background) mutant mice were used. We additionally used the more slowly degenerating *rd10* mice (C57Bl6/J) to assess primary rod degeneration at a longer timescale than in *rd1*, because in *rd10* retina, developmental and degenerative cell death are temporally less overlapping (Arango-Gonzalez et al., 2014; Sancho-Pelluz et al., 2008). Animals from all lines were used irrespective of gender. Wild-type and mutant *rd1* and *cpfl1* mice were crossbred with the transgenic mouse line *HR2.1:TN-XL* (C57BL/6J background), which expresses the Ca^2+^ biosensor TN-XL (Mank et al., 2006) under the control of the human red opsin promoter (HR2.1) selectively in cone photoreceptors (Wei et al., 2012). The mouse lines thereby generated were the *HR2.1:TN-XL* x *cpfl1* and *HR2.1:TN-XL* x *rd1* lines; for simplicity, we refer to these biosensor lines in the following as wt, *cpfl1,* and *rd1*. In the context of the present study, these mouse lines allowed for a direct identification of cones. Animals older than postnatal day 12 (P12) were sacrificed by CO_2_ asphyxiation followed by cervical dislocation. Mice younger than P12 were sacrificed by decapitation. All procedures were performed in accordance with the law on animal protection issued by the German Federal Government (Tierschutzgesetz) and approved by the institutional animal welfare office of the University of Tübingen.

### Genotyping for the rd8 mutation

Since the *rd8* mutation in the *Crb1* gene may constitute a significant confounding factor in studies on retinal degeneration (Mattapallil et al., 2012), all mouse lines used in this study were screened for the *rd8* mutation using a PCR amplification and Nde I restriction enzyme digestion. The PCR was run for 35 cycles using the forward primer 5’-GCCCCTGTTTGCATGGAGGAAACTTGGAAGACAGCTACAGTTCATAT-3’ and the reverse primer 5’-GCCCCATTTGCACACTGATGAC-3’ followed by Nde I digestion at 37°C over night. The PCR yielded a 244bp fragment for the *Crb1* wild-type gene and two fragments of 199bp and 45bp, respectively, for the *rd8* mutant allele. As positive controls, we used samples from heterozygous animals kindly provided by Dr. Ulrich F. Luhmann, University College London, UK. All animals used in our study were negative for the *rd8* mutation.

### Calpain activity assay

Following earlier work (Kulkarni et al., 2016), calpain activity was initially analysed in mice at P14, 18, 24, 30, 60, 90 and 120 (Table 1). In early degeneration *rd1* mouse and, for comparison, in wt animals, we added P10 and P12. Eyes were marked on the nasal side prior to enucleation, flash-frozen in liquid nitrogen, embedded in Tissue-Tek OCT compound (Sakura Finetek Europe, Alphen aan Den Rijn, Netherlands), and stored at −20°C until cryo-sectioning into 16 µm thick vertical sections. Sections were rehydrated in calpain reaction buffer (CRB; in g: 5.96 HEPES, 4.85 KCl, 0.47 MgCl_2_, 0.22 CaCl_2_ in 100ml ddH_2_O; pH 7.2) with 2 mM dithiothreitol (DTT), and then incubated for 2 hr at 37°C in CRB containing 50 µM of tBOC-Leu-Met-CMAC (Molecular Probes), a calpain-specific substrate, whose fluorescence increases after cleavage by calpain (Paquet-Durand et al., 2006).

**Table 1.**
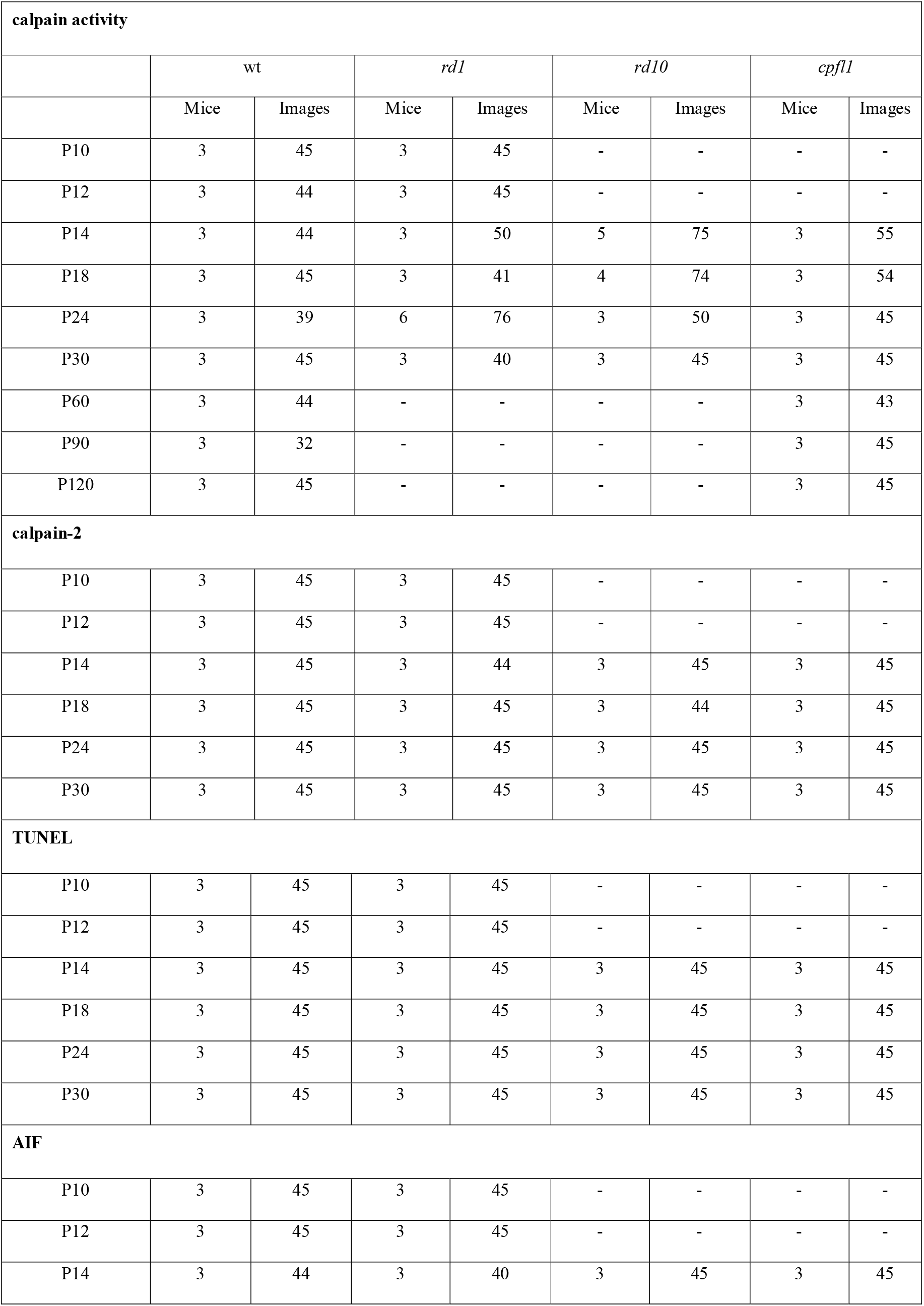
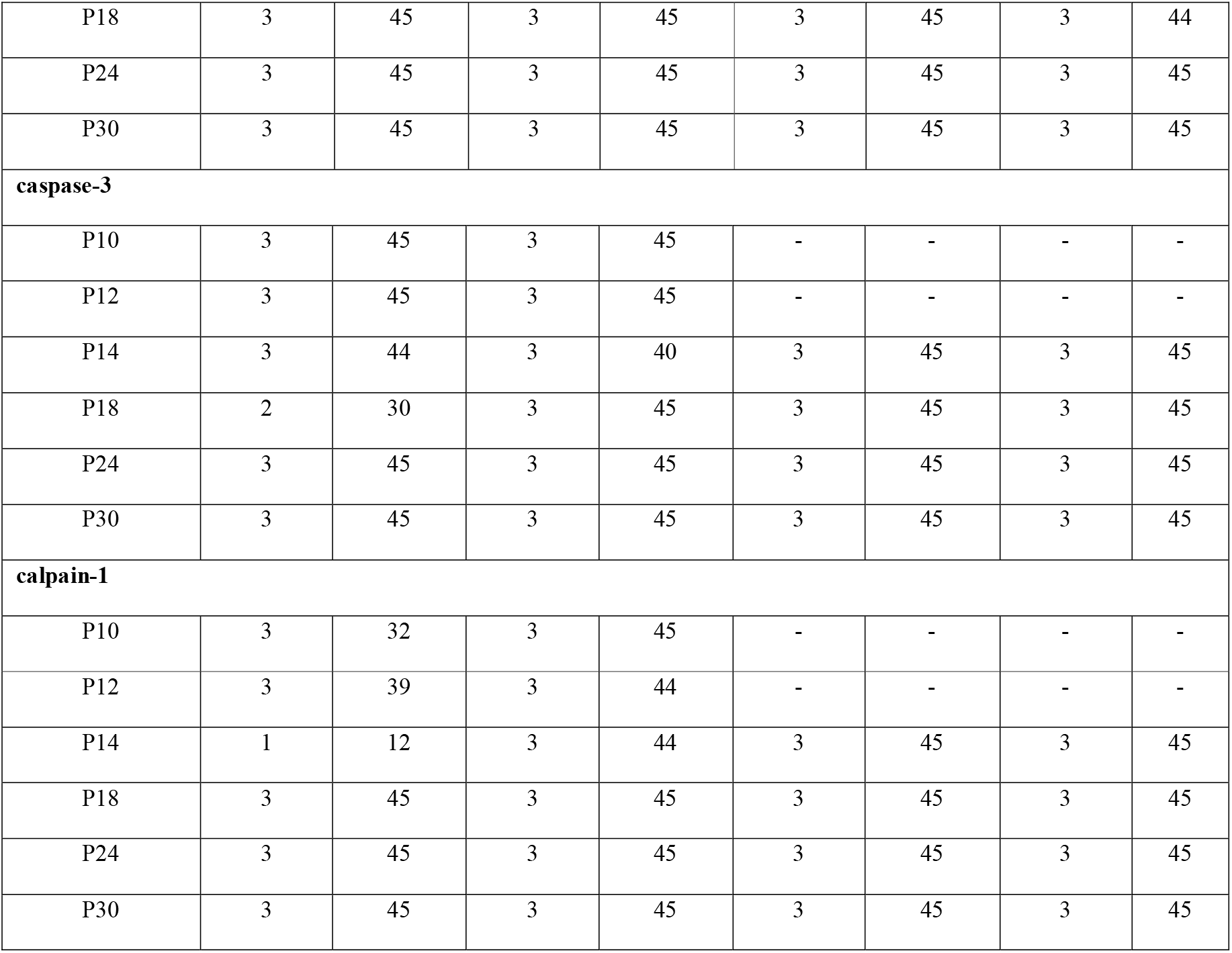
Number of mice used for each marker and number of images recorded. The table gives the information for all markers used, namely, calpain activity, calpain-1, calpain-2, TUNEL, caspase-3, AIF.

### Immunohistochemistry

We analysed mice at P10, 12, 14, 18, 24, and 30 (Table 1). Due to the lack of a discernible outer nuclear layer (ONL) in *rd1* and *rd10* animals after P30, and the lack of significant difference in calpain activity in *cpfl1* and wt post P30-animals, we limited our examination to time-points between P10 and P30. Eyes were fixed in 4% PFA in 0.1 M phosphate buffer saline (PBS, pH 7.4; for 45 min) and cryo-protected in sucrose gradients in PBS at room temperature for 2 hrs. Then, eyes were embedded in Tissue-Tek OCT compound (Sakura Finetek Europe, Alphen aan Den Rijn, Netherlands) and stored at −20°C until cryo-sectioning into 16 µm thick vertical sections. Sections were rehydrated with PBS, permeabilized in 0.3% Tween in PBS containing blocking solution (10% goat serum, 1% BSA). As primary antibodies, we used rabbit anti-calpain-2 (ab39165; 1:300; Abcam, Cambridge, UK), rabbit anti-calpain-1 (ab39170; 1:100; Abcam, Cambridge, UK), rabbit anti-cleaved-caspase-3 (#9664; 1:300, Cell Signalling Technology, Frankfurt, Germany: RRID: AB_2070042), and rabbit anti-AIF (ab1998; 1:350; Abcam, Cambridge, UK: RRID AB_302748). As secondary antibodies, we used goat anti-rabbit Alexa Fluor 488 (1:350; Molecular Probes, Eugene, USA: RRID AB_143165).

### Cell death detection

Mice were analysed at P10, 12, 14, 18, 24, and 30 (Table 1). The terminal deoxynucleotidyl transferase dUTP nick end labelling (TUNEL) assay was performed using the TMR red kit from Roche Diagnostics (Mannheim, Germany). Sections were incubated in blocking solution (1% BSA, 10% normal goat serum, 1% fish gelatine) for 1 hr after 5 min incubation with alcohol acetic acid mixture (62% EtOH, 11% Acetic Acid, 27% H_2_O). Sections were then stained using TMR red TUNEL kit as per manufacturer’s instructions.

### Microscopy and image processing

Z-stack images were captured on an Imager Z1 ApoTome microscope using a 20x air objective (0.8 NA; *cf*.), and Zen (v.2.3 Pro) software (Carl Zeiss Microscopy GmbH, Oberkochen, Germany). Zen Lite v.2.3 software was used to reconstruct images and ImageJ 1.52e (http://imagej.nih.gov/ij) was used to count positive markers in each image. Figures were generated in IGOR Pro (Wavemetrics, Lake Oswego, USA) and arranged in Canvas 11 (ACD Systems International Inc., Seattle, USA).

### Analysis of marker data

Data was obtained from at least three different animals for each parameter examined. For each animal, vertical sections were quantified, using collapsed Z-stacks of n= 10-22 images, spaced at 0.75 µm z-distance, acquired at 20x magnification (Fig. 1). For each section, five regions across the retina were imaged and analysed. Here, the central retina was taken as the medial portion along both the dorso-ventral and naso-temporal axis, as well as, dorsomedial and ventromedial sections close to the optic nerve. Calpain-active, as well as calpain-2-, calpain-1-, caspase-3-, AIF-, and TUNEL-labelled cells were counted and expressed as number of positive cells in the ONL per 1,000 µm^2^ in all mouse lines, and at all time-points. Statistical comparisons were made using the Kruskal–Wallis one-way analysis of variance test using GraphPad Prism 7 (La Jolla, USA). Statistical comparisons were confined to different mouse lines of the same age and not across ages.

**Figure 1.**
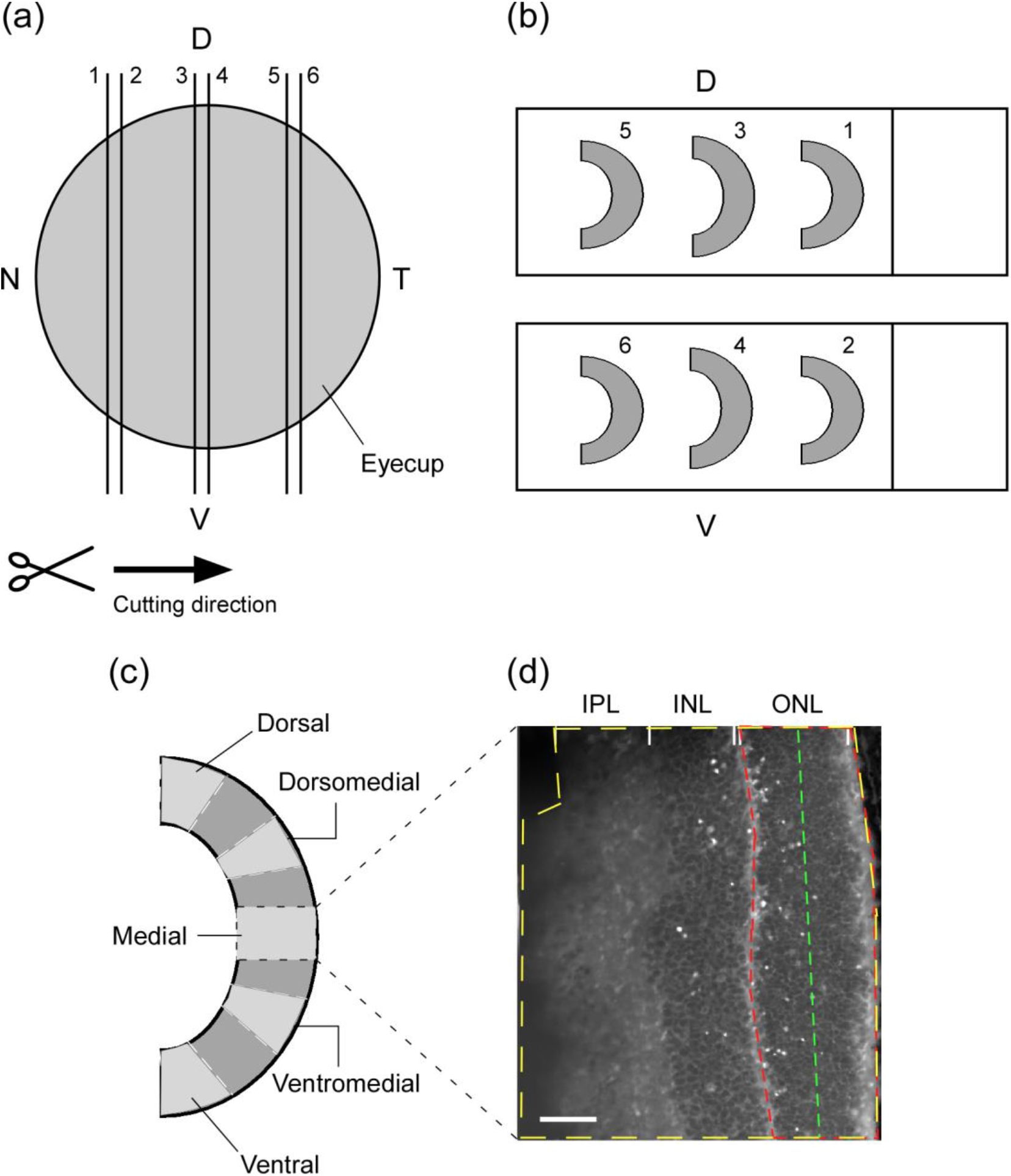
Illustration of the preparation protocol. Mouse eyes were marked nasally before being enucleated, dissected, fixed, and flash frozen in liquid nitrogen before sectioning. Eyes were sectioned from the nasal side towards the temporal side along the dorso-ventral axis (**A**). Consecutive sections were kept on consecutive slides (**B**). Sections were imaged along the dorso-ventral axis at five locations: Dorsal, Dorsomedial, Medial, Ventromedial and Ventral (**C**). Images were then analysed (**D**) by measuring the whole retinal area imaged (yellow dashed line), the ONL area imaged (red dashed line) and, ONL length imaged (green dashed line) for each image. ONL width was calculated by dividing the area by the length for each image. Scale bar: 50 μm.

### Gaussian process modelling

Gaussian Process models (GPs) (Williams & Rasmussen, 2006) were used to estimate the mean and standard deviation of the activity of each cell death marker, for each mouse line, at the observed time-points. These models were used as they are able to accommodate the non-linear change in the level of each marker, and provide a generative model for estimating properties of interest (in our case, the likely peaks for each marker). Modelling was performed in Python 3.5 using the GPy (GPy, 2014) library.

Prior to model fitting, a square root transformation was applied to the observed activity of each marker to accommodate the left-skew and zero-bound; the inverse transformation was applied for subsequent inference. The GPs inferred how the observed activity of each marker, *y*, varied over time, *x*, for each mouse line. The GP (Eq. 1) was defined by the mean activity function *µ*(*x*), the covariance of this activity *K*(*x*, *x^’^*) (Eq. 2), and additive zero-mean Gaussian observation noise *ε* (Eq. 3). A radial basis function kernel was used to model the covariance of the signal over time. The model was parameterised by the signal variance 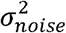, length scale *l*, and noise variance 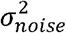; these parameters were inferred using noise noise the L-BFGS-B maximum likelihood algorithm.

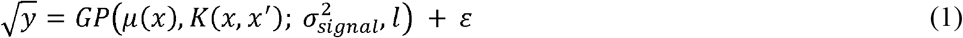

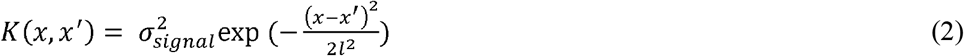

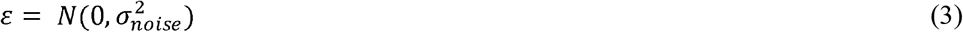

Once fitted, the GPs could be used as generative models to compute bootstrap estimates of molecular sequences. We drew 10,000 samples from each model and identified the maxima in each sample. The marginal distribution of these maxima was used as an estimate of the timing of the peak of activity in each model. For each of the samples, we also identified the relative ordering of the peaks of each molecular marker, to derive an approximate molecular sequence. The probability of any given molecular sequence was calculated as the number of times it was present in these samples, as a proportion of the total number of samples which were drawn from the models. We excluded those sequences that had a probability of 0.05 or lower.

### Generation of heat maps

During retinal sectioning, order and orientation of sections were maintained (Fig. 1). Hence, the retinal location of each image could be determined using the section number and the distance from the optic nerve head along the section, providing x- (nasal-temporal) and y-coordinates (dorsal-ventral), respectively. To visualise the spatio-temporal progression of the markers, the mean activity at each location along the dorsal-ventral axis was used (averaging over the nasal-temporal dimension) and plotted against each time-point. Heat-maps were generated in Python 3 using the matplotlib library.

## 3. Results

To link Ca^2+^ dysregulation to photoreceptor cell death, we utilised RD mouse models with mutations in *Pde6* genes. The *rd1* and *rd10* lines carry different mutations in the *Pde6b* gene, whereas the *cpfl1* line suffers from a mutation in the homologous, cone-specific *Pde6c* gene. To align our experiments with earlier studies of mouse photoreceptor degeneration, we first compared changes in outer nuclear layer (ONL) thickness over the first post-natal month between the mutants and wt mice (Fig. 2). The number of photoreceptor rows dramatically decreased in post-P12 *rd1* and in *rd10* after P18, illustrating the difference in onset of photoreceptor cell death in these models (Arango-Gonzalez et al., 2014). The *rd10* degeneration is less aggressive than *rd1* as it takes longer for complete ablation of the ONL in *rd10* than in *rd1*. In contrast, in the cone degeneration *cpfl1* mutant the ONL thickness remains similar to wt, because in mice, rods outnumber cones by a factor of 25 to 50 (Jeon, Strettoi, & Masland, 1998, Behrens et al., 2016), and primary cone loss does not produce secondary rod loss.

**Figure 2.**
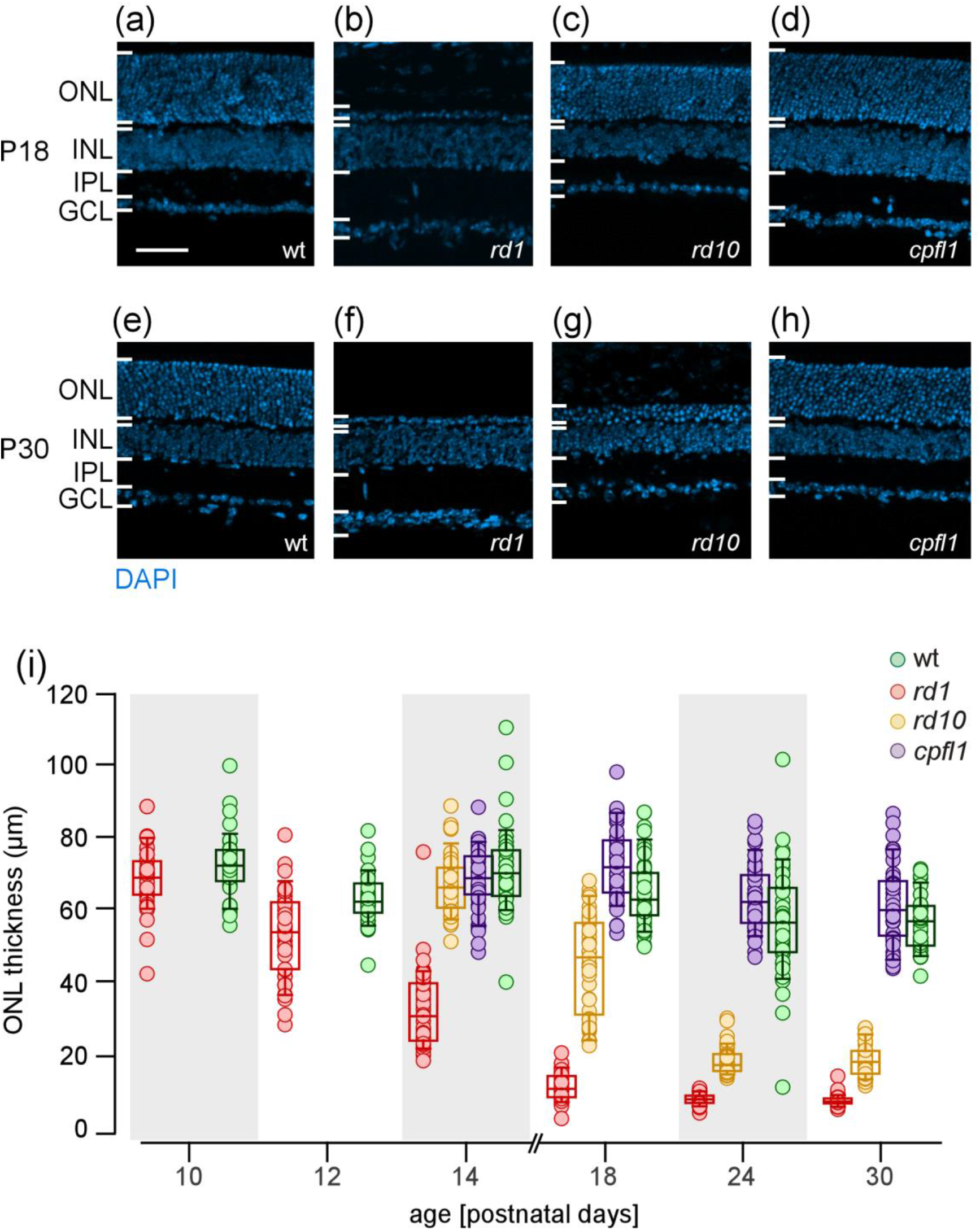
Photoreceptor degeneration-related cell loss in different mouse lines. **A-H**, representative images of sections from *wt* (**A**, **E**), *rd1* (**B**, **F**), *rd10* (**C**, **G**), and *cpfl1* (**D**, **H**) analysed based DAPI-staining, images at P18 (top) and P30 (bottom) of each mouse line. **I**, cell loss was measured using outer nuclear layer (ONL) thickness during the first post-natal month in *rd1* (cherry), *rd10* (amber), *cpfl1* (lilac), and wild-type (wt, emerald) mice (n=3 animals per mouse line and time-point).

### Calpain activity increases in ONL of degenerating retina

We next aimed at elucidating the role of Ca^2+^ dysregulation in photoreceptor cell death. To this end, we focussed first on the activity and regulation of Ca^2+^-dependent calpain-type proteases, which can be considered as surrogate markers for Ca^2^ dysregulation (Croall & Ersfeld, 2007; Goll et al., 2003). In *rd1* retina, the number of ONL cells showing increased calpain activity was significantly higher compared to wt and *cpfl1*, at all time-points, with a peak of activity at P12 (Fig. 3A, D; for detailed statistics of all comparisons, see Table 2). Calpain activity in *rd10* ONL was also significantly increased compared to wt and *cpfl1* (at P18, 24, and 30), peaking around P18 (Fig. 3B, D). In c*pfl1*, calpain activity tended to be higher than in wt, yet, without a clear peak within the inspected time frame (Fig. 3C, D). Overall, *rd1* retinae displayed the highest levels of calpain activity. These data are in line with other studies (Arango-Gonzalez et al., 2014; Paquet-Durand et al., 2006) that showed increases in calpain activity at specific time-points in models of RP and achromatopsia.

**Figure 3.**
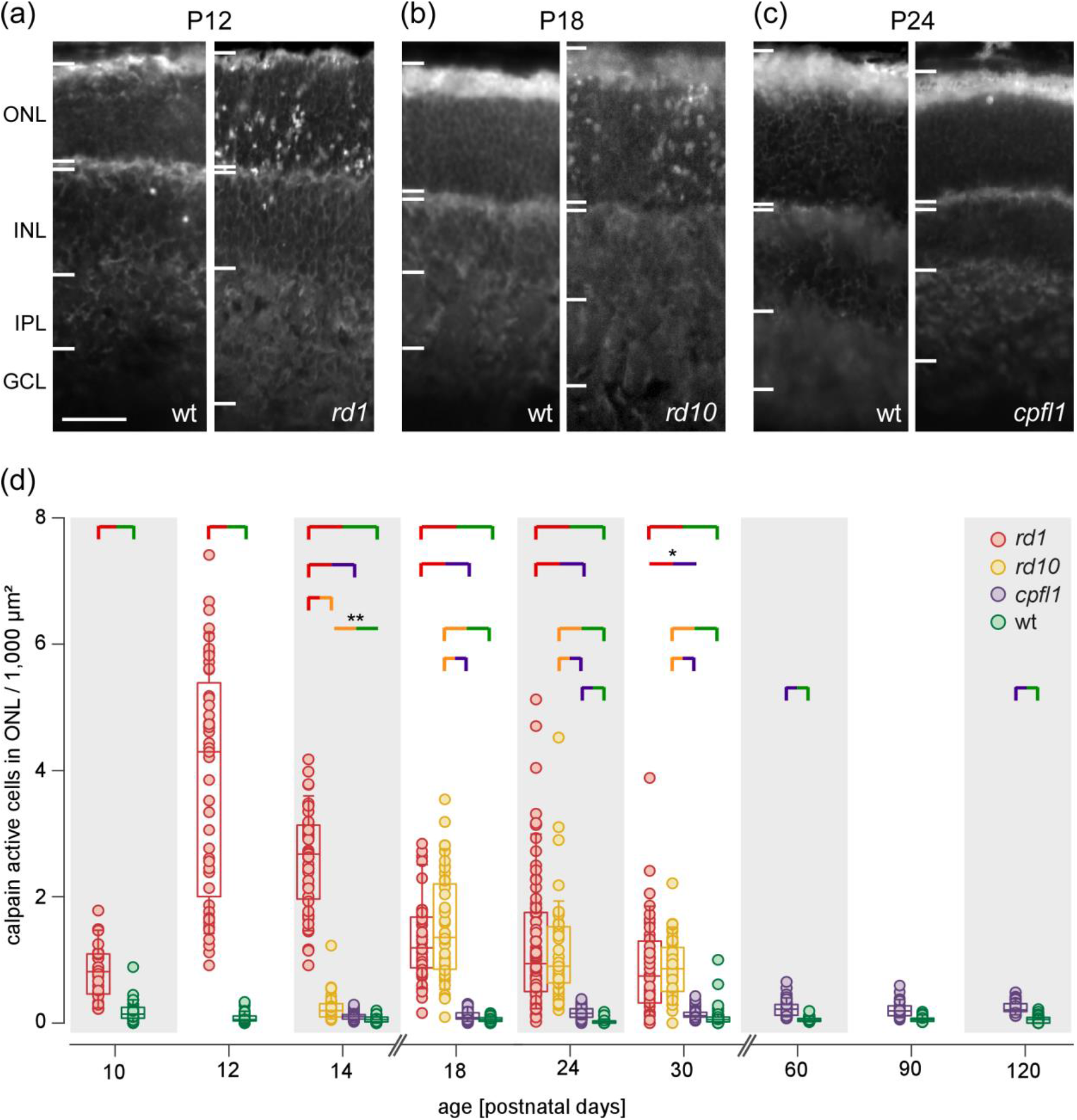
Temporal progression of calpain activity in different mouse lines. **A-C**, representative images of sections from *rd1* (**A**), *rd10* (**B**), and *cpfl1* (**C**) analysed with the calpain assay; each image of a mutant retina (left) is age-matched with a wt control (right). **D**, normalised number of fluorescent cells in the ONL (per 1,000 μm^2^), indicative of calpain activity in photoreceptors, as a function of age (n=45 observations obtained on 3 animals per mouse line and time-point). Statistical analysis was Kruskal–Wallis one-way analysis of variance, brackets denote statistical significance (p < 0.001), unless otherwise indicated (**; p<0.01, *; p<0.05). Scale bar: A-C, 50 μm.

**Table 2:** Values and statistics from each marker recorded. Median values plus/minus the standard error of the mean recorded from each mouse model at each time-point and for each marker.

| **calpain activity** |  |  |  |  |
| --- | --- | --- | --- | --- |
|  | *Wild type* | *rd1* | *rd10* | *cpfl1* |
| P10 | 0.14±0.023 | 0.78±0.068 | - | - |
| P12 | 0.055±0.012 | 4.3±0.27 | - | - |
| P14 | 0.053±0.0074 | 2.69±0.11 | 0.11±0.019 | 0.09±0.008 |
| P18 | 0.051±0.0057 | 1.19±0.13 | 1.36±0.145 | 0.08±0.01 |
| P24 | 0.0255±0.006 | 1.001±0.13 | 0.878±0.122 | 0.15±0.01 |
| P30 | 0.06±0.02 | 0.74±0.12 | 0.82±0.068 | 0.12±0.01 |
| P60 | 0.044±0.005 | - | - | 0.22±0.02 |
| P90 | 0.054±0.007 | - | - | 0.176±0.02 |
| P120 | 0.058±0.008 | - | - | 0.21±0.016 |
| **TUNEL** |  |  |  |  |
| P10 | 0.046±0.01 | 1.15±0.116 | - | - |
| P12 | 0.078±0.016 | 6.1±0.42 | - | - |
| P14 | 0.156±0.011 | 3.63±0.18 | 0.17±0.12 | 0.16±0.017 |
| P18 | 0.14±0.011 | 2.94±0.3 | 1.76±0.23 | 0.21±0.019 |
| P24 | 0.086±0.013 | 1.44±0.14 | 2.7±0.16 | 0.19±0.01 |
| P30 | 0.03±0.0088 | 1.40±0.101 | 1.08±0.08 | 0.1±0.02 |
| **calpain-2** |  |  |  |  |
| P10 | 0.034±0.006 | 0.46±0.05 | - | - |
| P12 | 0±0.006 | 0.95±0.14 | - | - |
| P14 | 0.026±0.006 | 1.10±0.13 | 0±0.01 | 0.016±0.0049 |
| P18 | 0.019±0.005 | 0.3±0.033 | 0.41±0.09 | 0±0.003 |
| P24 | 0±0.007 | 0.26±0.07 | 0.29±0.05 | 0.024±0.0038 |
| P30 | 0±0.003 | 0.19±0.088 | 0.73±0.1 | 0±0.004 |
| **calpain-1** |  |  |  |  |
| P10 | 0±0 | 0±0 | - | - |
| P12 | 0±0.0035 | 0±0.0015 | - | - |
| P14 | 0±0 | 0±0.002 | 0±0.002 | 0±0.0007 |
| P18 | 0±0.0007 | 0±0.0016 | 0±0.04 | 0±0.002 |
| P24 | 0±0.0004 | 0±0.004 | 0±0.025 | 0±0.0009 |
| P30 | 0±0.002 | 0±0.014 | 0.38±0.05 | 0±0.0009 |
| **caspase-3** |  |  |  |  |
| P10 | 0±0.00095 | 0±0 | - | - |
| P12 | 0±0.00057 | 0±0 | - | - |
| P14 | 0±0 | 0±0.004 | 0±0 | 0±0.0009 |
| P18 | 0±0 | 0±0.019 | 0±0.006 | 0±0.0004 |
| P24 | 0±0 | 0±0.045 | 0±0.007 | 0±0 |
| P30 | 0±0 | 0±0.008 | 0±0.001 | 0±0.0004 |
| **AIF** |  |  |  |  |
| P10 | 0±0.0006 | 0±0.002 | - | - |
| P12 | 0±0.001 | 0±0.0025 | - | - |
| P14 | 0±0.003 | 0±0.0035 | 0±0.003 | 0±0.004 |
| P18 | 0±0.001 | 0±0.018 | 0.45±0.1 | 0±0.004 |
| P24 | 0±0.0007 | 0±0.027 | 0.91±0.091 | 0±0.0003 |
| P30 | 0±0.0023 | 0±0.085 | 0.65±0.08 | 0±0.0008 |

### Activation of calpain-2, but not calpain-1, increases during disease progression

We then investigated whether the high levels of calpain activity observed in the ONL could be attributed to specific calpain isoforms. We focussed on calpain-1 and calpain-2, which have been reported to be ubiquitously expressed in all mammalian cells and to play opposite roles in neurodegeneration (Chen et al., 2007; Goñi-Oliver et al., 2007). We hypothesized that calpain-2 may contribute to the observed increased calpain activity, as it requires a high [Ca^2+^] that is considered beyond the physiological range of photoreceptors (Goll, 1995). Using antibodies recognizing the activated proteases, we found significantly increased numbers of calpain-2 positive cells in both *rd1* and *rd10* ONL, when compared to *cpfl1*, as well as to wt controls (Fig. 4A, B, D; for all statistics, see Table 2). The peaks appeared to coincide with those for the calpain assay, namely at P12 (*rd1*) and P18 (*rd10*). In *cpfl1*, the number of calpain-2 positive cells was slightly increased over the wt level (Fig. 4C, D), however, without a clear peak. To our surprise, a significant increase in calpain-1 positive cell numbers from wt level was observed in *rd10* (Fig. 5A-C). Calpain-1 labelling increased over background levels around P18 and was even more pronounced at the end of the observed time window (P30) (Fig. 5D). Next to *rd10*, only *rd1* showed a detectable (but not significant) increase in calpain-1 positive cell counts.

**Figure 4.**
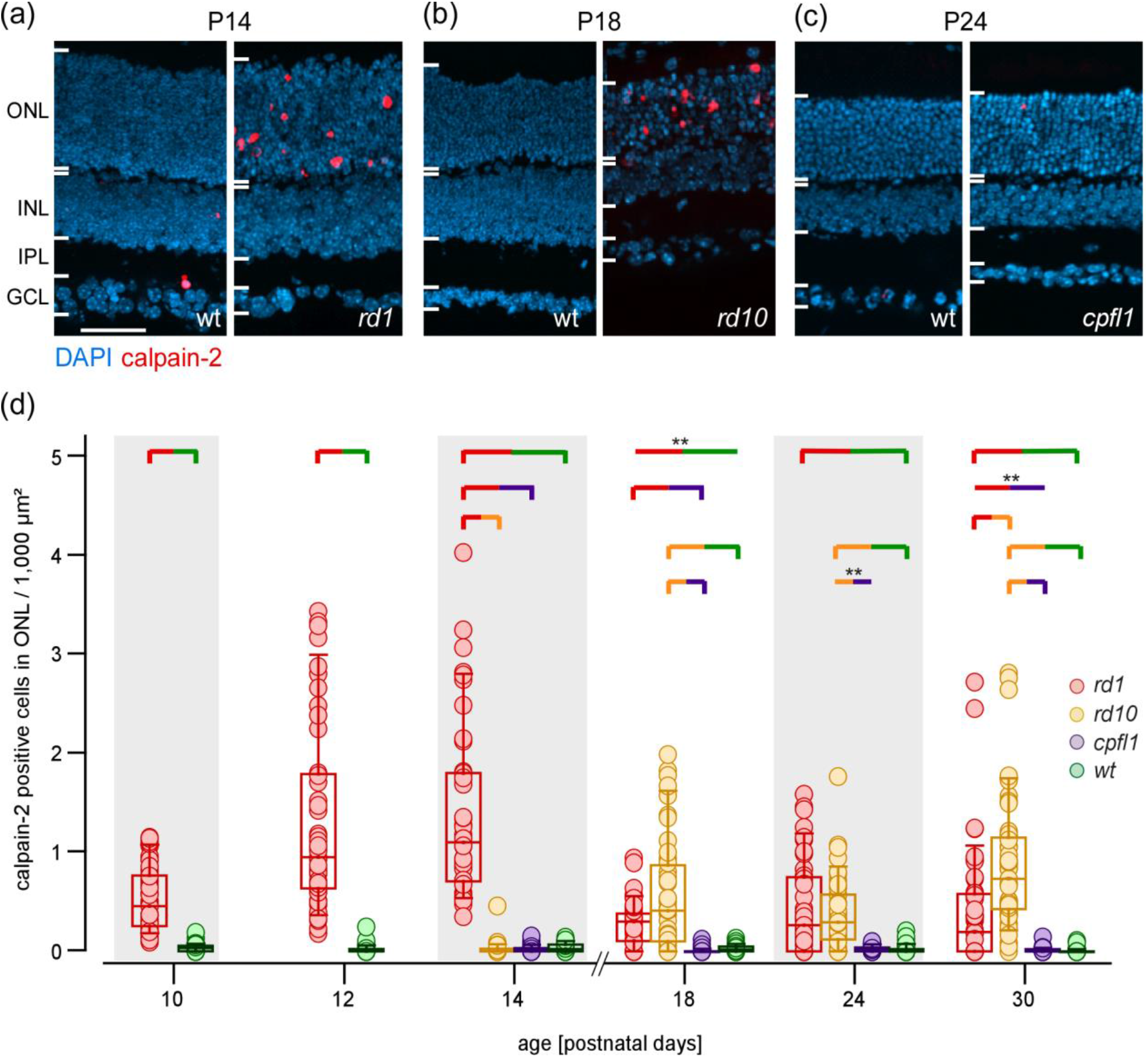
Calpain-2 becomes activated during photoreceptor degeneration. **A-C**, representative images of sections from *rd1* (**A**), *rd10* (**B**), and *cpfl1* (**C**) analysed for calpain-2 immuno-reactivity; each image of a mutant retina (left) is age-matched with a wt control (right). **D**, normalised number of fluorescent cells in the ONL (per 1,000 μm^2^), indicative of calpain-2 immunoreactive photoreceptors, as a function of age (n=45 observations obtained on 3 animals per mouse line and time-point). Statistical analysis was Kruskal-Wallis one-way analysis of variance, brackets denote statistical significance (p < 0.001), unless otherwise indicated (**; p<0.01). Scale bar: A-C, 50 μm

**Figure 5.**
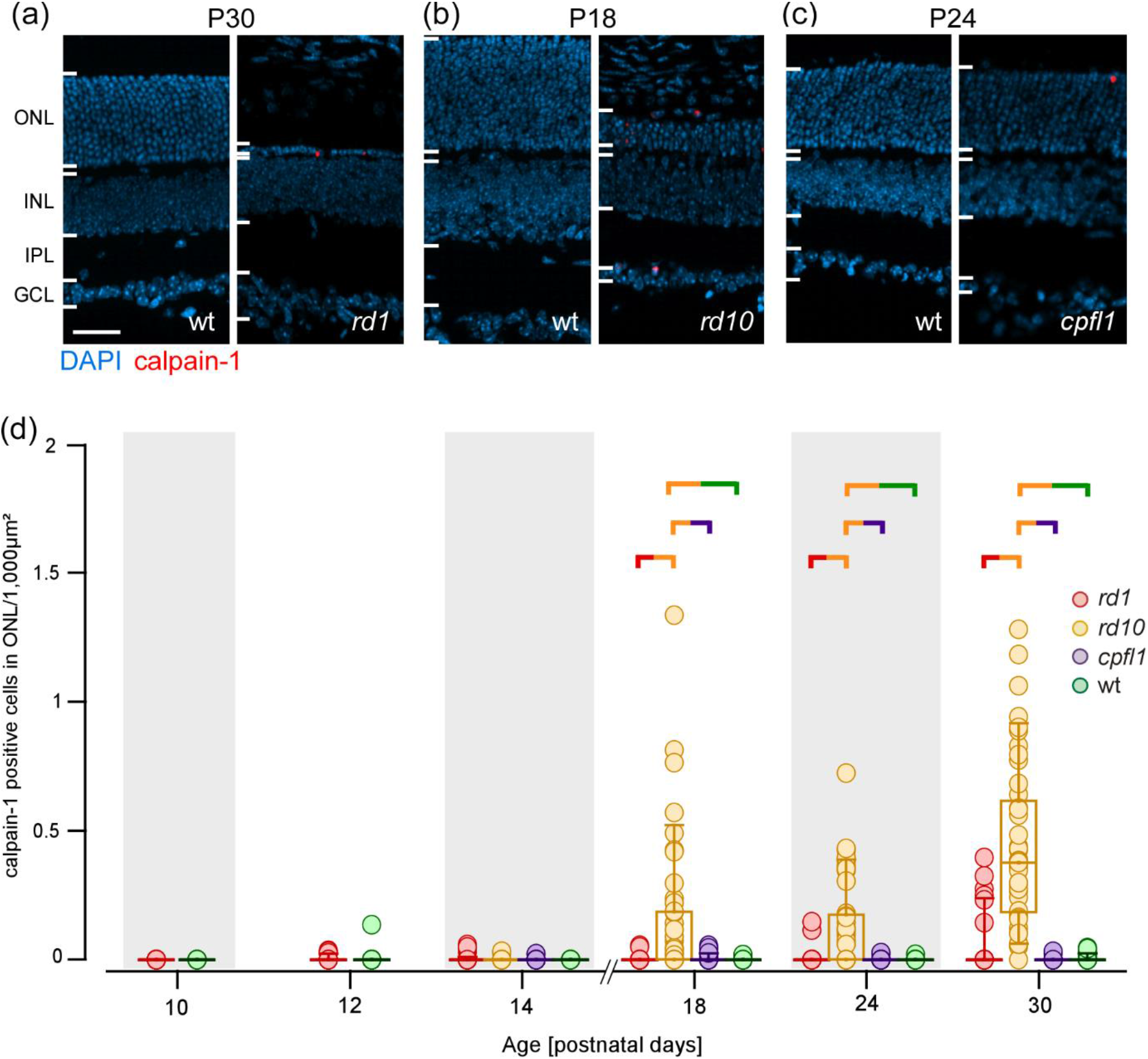
Calpain-1 immunolabeling in different RD and wt mouse lines. **A-C**, representative images of sections from *rd1* (**A**), *rd10* (**B**), and *cpfl1* (**C**) analysed for calpain-1 positivity; each image of a mutant retina (left) is age-matched with a wt control (right). **D**, normalised number of fluorescent cells in the ONL (per 1,000 μm^2^), indicative of calpain-1 immunoreactive photoreceptors, as a function of age. Statistical analysis was performed using Kruskal-Wallis one-way analysis of variance, brackets denote statistical

Taken together, we found differential patterns of calpain isoform activation: While calpain-2 was strongly activated in *rd1* and *rd10* (and much less so in *cpfl1*), calpain-1 was more strongly activated in *rd10* compared to the other lines. Moreover, the calpain-2 expression peak coincides with the peak of calpain activity assay data and precedes that of calpain-1.

### Cell death coincides with calpain activity

To compare the activity and expression of calpain in the ONL with the incidence of cell death, we used the TUNEL assay, which labels nick-ends in a cell’s DNA. While TUNEL is an excellent marker for cell death, for the present study it is important to mention that TUNEL does not discriminate between different cell death mechanism(s) (Kraupp et al., 1995). In agreement with earlier work (Arango-Gonzalez et al., 2014), we found significantly increased numbers of TUNEL-positive cells in *rd1* and *rd10* ONL (Fig. 6A, B, D: for all statistics, see Table 2). A slight increase in TUNEL-positive cell number was also seen in the *cpfl1* mouse (Fig. 6C, D) albeit without discernible peak. In *rd1* and *rd10* animals, the peaks in TUNEL-positive ONL cell numbers (P12 in *rd1*; P24 in *rd10*) coincided temporally with those of calpain activity and calpain-2 labelling.

**Figure 6.**
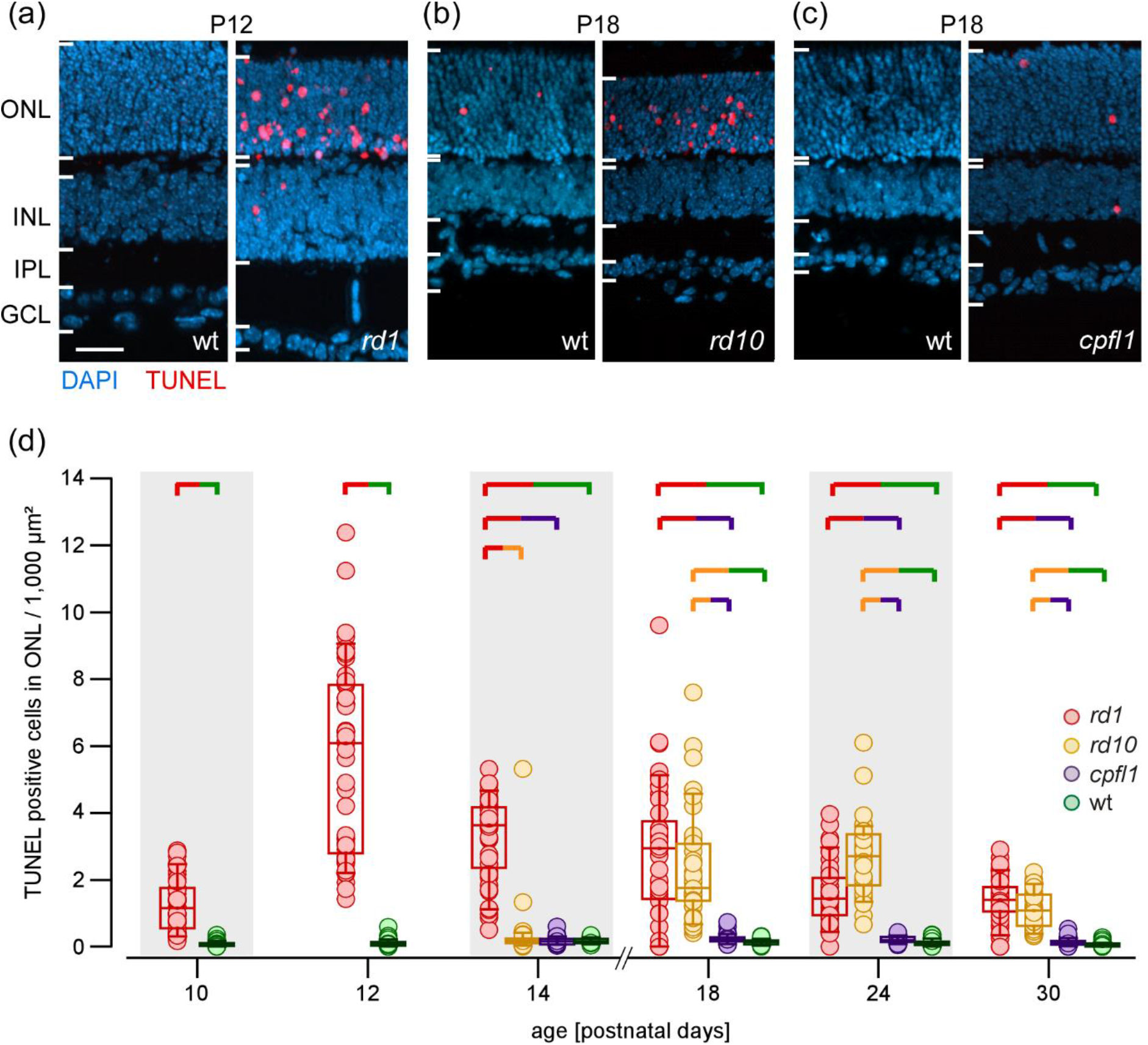
Temporal progression of cell death in different RD and wt mouse lines. **A-C**, representative images of sections from *rd1* (**A**), *rd10* (**B**), and *cpfl1* (**C**) analysed with the TUNEL assay; each image of a mutant retina (left) is age-matched with a wt control (right). **D**, normalised number of fluorescent cells in the ONL (per 1,000 μm^2^), indicative of TUNEL positivity in photoreceptors, as a function of age (n=45 observations obtained on 3 animals per mouse line and time-point). Statistical analysis was Kruskal-Wallis one-way analysis of variance, brackets denote statistical significance of *** ≤ 0.001 unless otherwise stated. Scale bar: A-C, 50 μm.

### AIF and caspase-3 levels argue against classical apoptosis in dying rods

How high intracellular [Ca^2+^] and Ca^2^-dysregulation are linked to photoreceptor cell death is still unclear. Prominent candidates for Ca^2+^-dependent downstream pathways are apoptotic cell death through caspase effectors (Orrenius et al., 2003) and non-apoptotic cell death involving calpain activation (Arango-Gonzalez et al., 2014; Doonan et al., 2005). To distinguish between these two proposed pathways, we used antibodies for cleaved caspase-3, a marker protein for apoptosis (Mazumder, Plesca, & Almasan, 2008), and AIF, a marker for programmed necrosis (Arango-Gonzalez et al., 2014; Wang et al., 2018).

AIF-positive cells were observed in the ONL of all disease models (Fig. 7A-D). Significant increases in AIF-positive cell numbers in *rd10* occurred at the same time-point (P18; for all statistics, see Table 2) as those of TUNEL and calpain activity. In *rd1* retina, a significant increase in AIF-positive cell numbers was seen only at the P30 time-point. In addition, a small, non-significant increase in AIF-positive cells number was seen in *cpfl1* retina.

**Figure 7.**
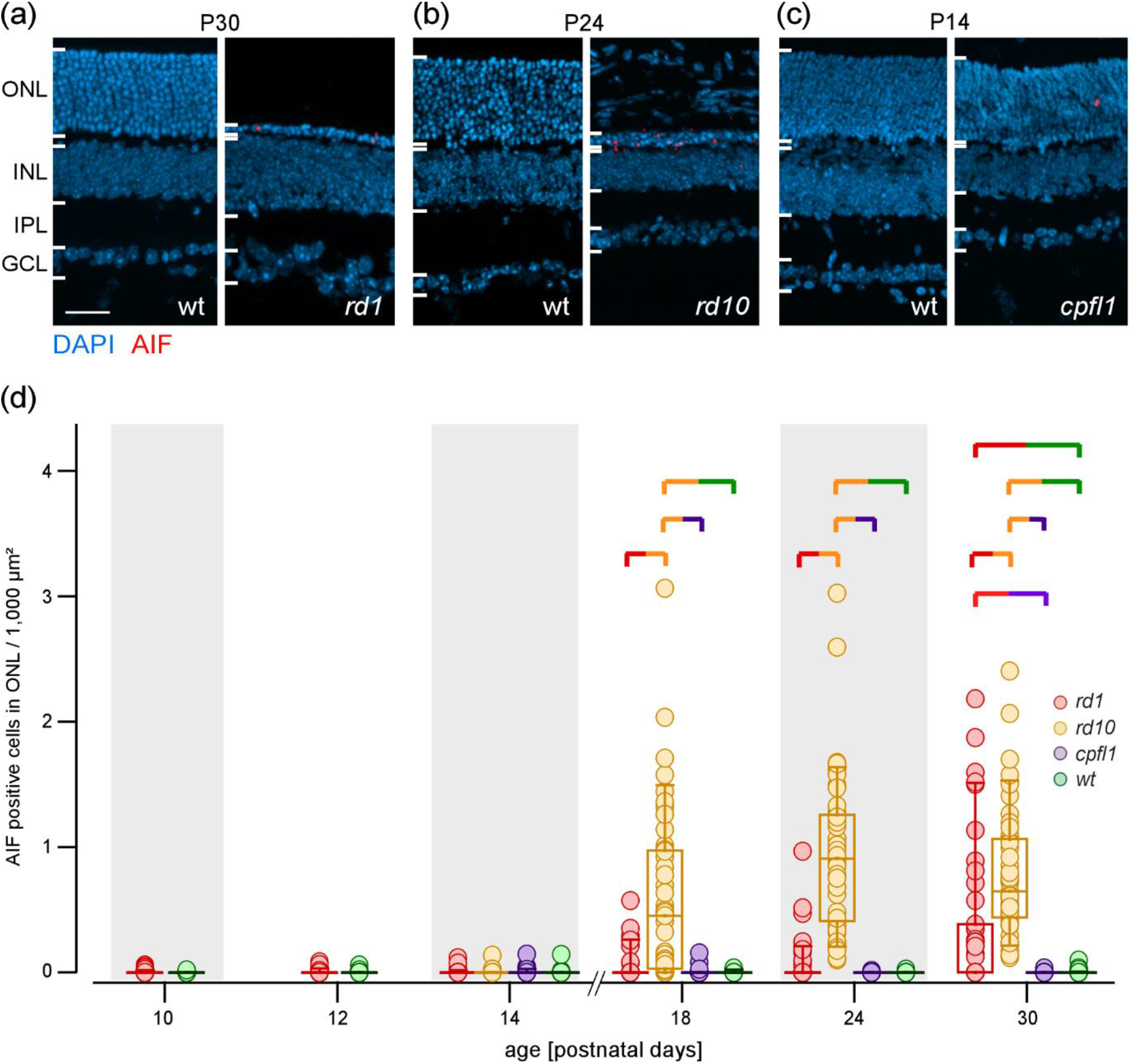
AIF immunolabeling in different RD and wt mouse lines. **A-C**, representative images of sections from *rd1* (**A**), *rd10* (**B**), and *cpfl1* (**C**) analysed for AIF immunofluorescence; each image of a mutant retina (left) is age-matched with a wt control (right). **D**, normalised number of fluorescent cells in the ONL (per 1,000 μm^2^), indicative of AIF immunoreactive photo-receptors, as a function of age (n=45 observations obtained on 3 animals per mouse line and time-point). Statistical analysis was Kruskal-Wallis one-way analysis of variance, brackets denote statistical significance of*** ≤ 0.001 unless otherwise stated. Scale bar: A-C, 50 μm.

Increases in the numbers of cells showing caspase-3 activation were observed in all mouse lines (Fig. 8A-C), though to a low level (Fig. 8D) and not at all time-points (for all statistics, see Table 2). Significantly higher numbers of caspase-3 positive cells were seen in *rd1* retina from P18 on, peaking at P24. Caspase-3 positive cells were often overlapping with cones, which were marked by the TN-XL biosensor expression (Wei et al., 2012). Indeed, when we analysed for *rd1* cones expressing caspase-3, we found that at P24 a substantial number of cones were positive for activated caspase-3 (Fig. 9A-D). This delayed caspase-3 activation is compatible with the execution of apoptosis during secondary *rd1* cone degeneration.

**Figure 8.**
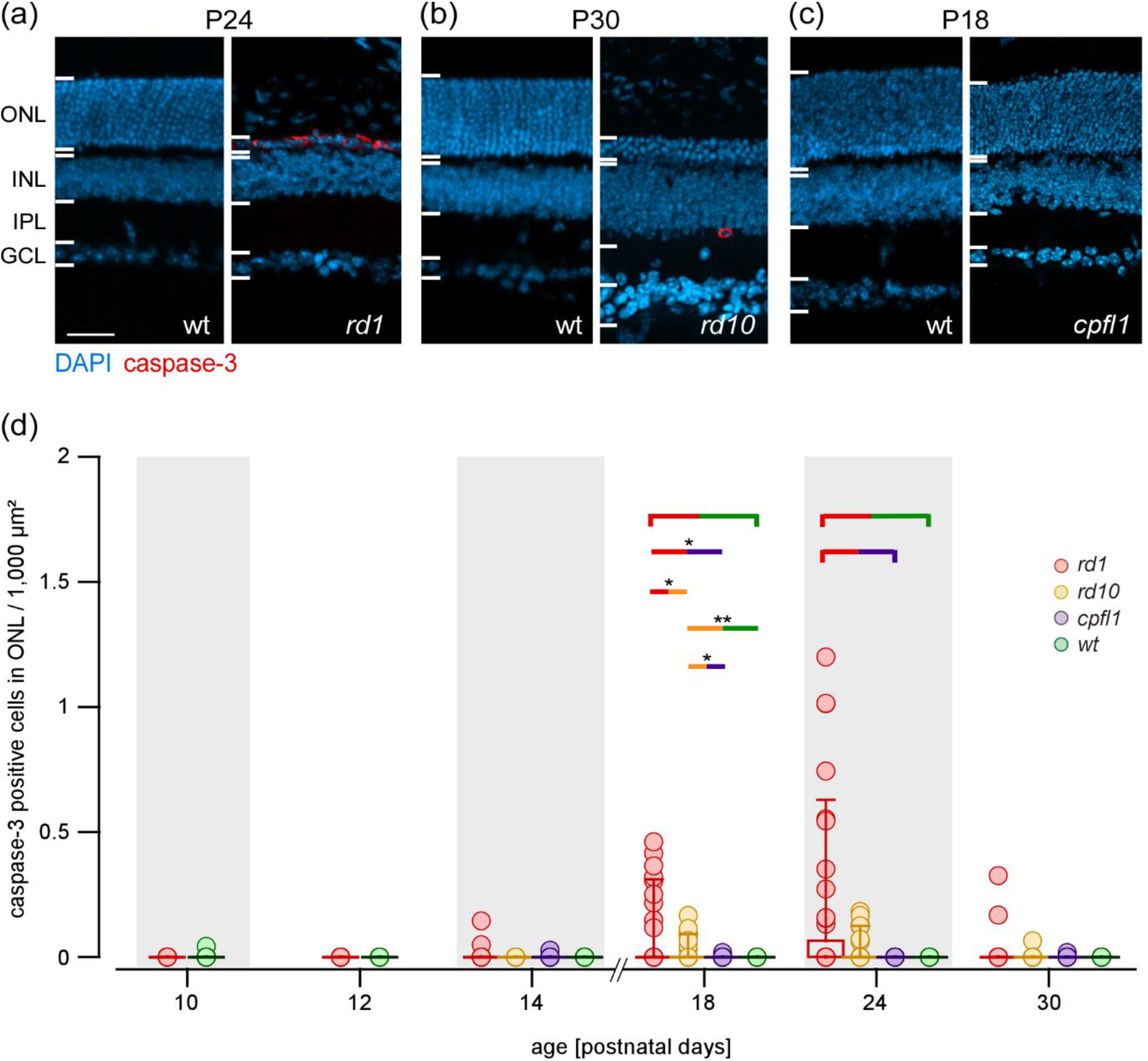
Activated caspase-3 in wt and RD mouse models. **A-C**, representative images of sections from *rd1* (**A**), *rd10* (**B**), and *cpfl1* (**C**) stained with an antibody against activated caspase-3; each image of a mutant retina (left) is age-matched with a wt control (right). **D**, normalised number of fluorescent cells in the ONL (per 1,000 μm^2^), indicative of photo-receptors with activated caspase-3, as a function of age (n=45 observations obtained on 3 animals per mouse line and time-point). Statistical analysis was Kruskal-Wallis one-way analysis of variance, brackets denote statistical significance of*** ≤ 0.001 unless otherwise indicated (**; p <0.01, *; p<0.05). Scale bar: A-C, 50 μm.

**Figure 9.**
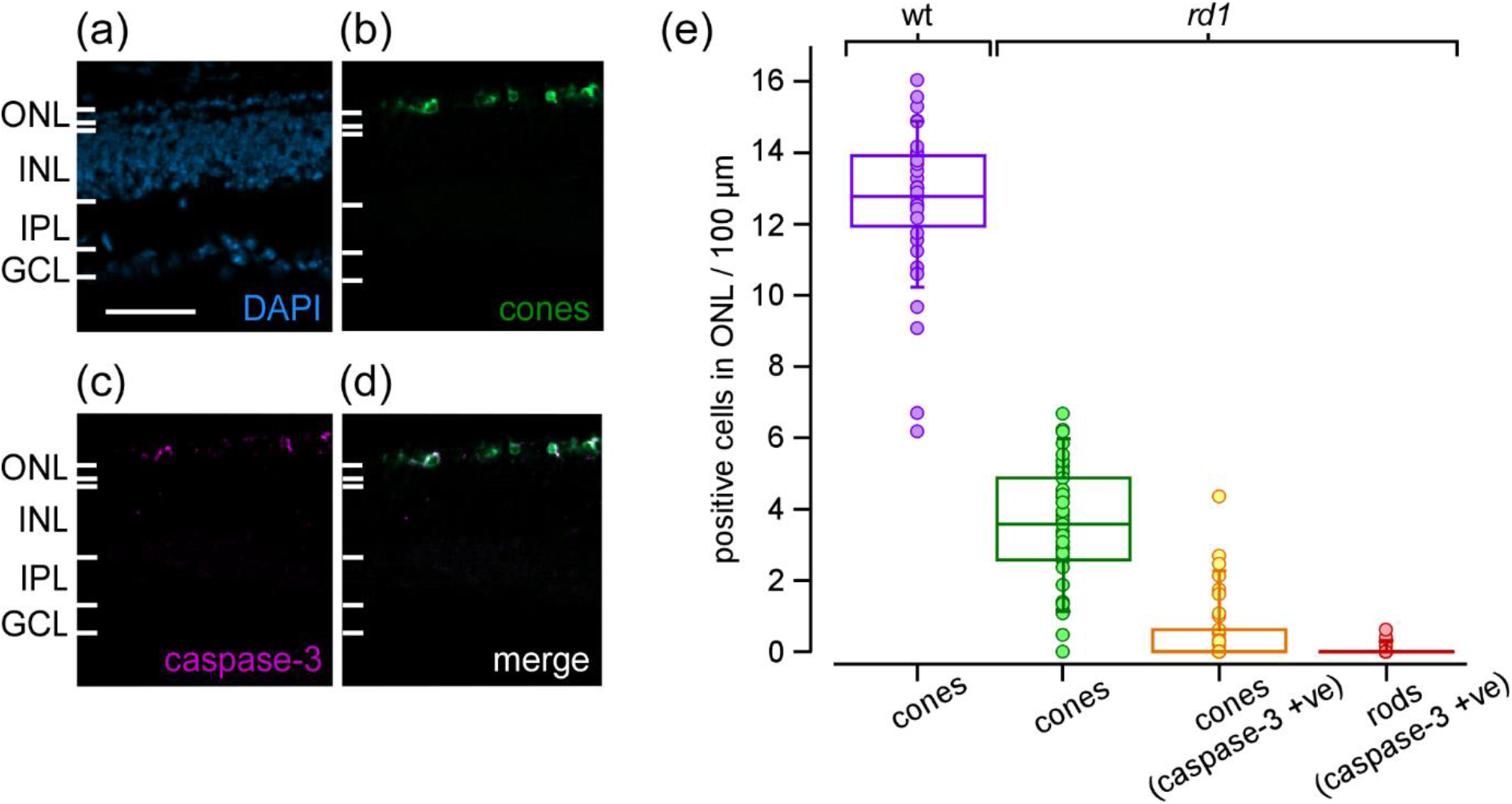
Activated caspase-3 in degenerating cone photoreceptors. **A-D**, representative images of sections from *rd1* P24 showing DAPI (**A**), cones expressing the TN-XL biosensor (**B**), activated caspase-3 (**C**), and the merged image (**D**). **E**, normalised number of fluorescent cells in the ONL (per 100 μm), indicative of cones in the wt retina at P24 (lilac), cones not showing caspase-3 activation (green), cones positive for activated caspase-3 (caspase-3 positive (+ve); yellow) and, rods positive for activated caspase-3 (caspase-3 +ve; red; n=45 observations obtained on 3 animals per mouse line and time-point). In total 778 cones were counted in 45 images, with 111 cones (14%) positive for caspase-3. Scale bar: A-D, 50 μm.

### Spatio-temporal mapping of degenerative markers in the retina

In the experiments so far, we noticed that the distribution of data points for a certain time-point was often quite broad and featured multiple peaks (*e.g.* Fig. 3D, *rd1* at P12; Fig. 4D, *rd10* at P24). Possible explanations for this are variability between individual mice and variability of degeneration state across the retina. We tested the latter by resolving the distributions of the markers along the retina’s dorso-ventral axis for all time-points and lines (Fig. 10). To this end, we collapsed data points along the naso-temporal axis, calculating the mean distribution along the dorso-ventral axis for each time-point, and visualized the whole time series as spatio-temporal heat maps (for original maps, see Figs. 11-16).

**Figure 10.**
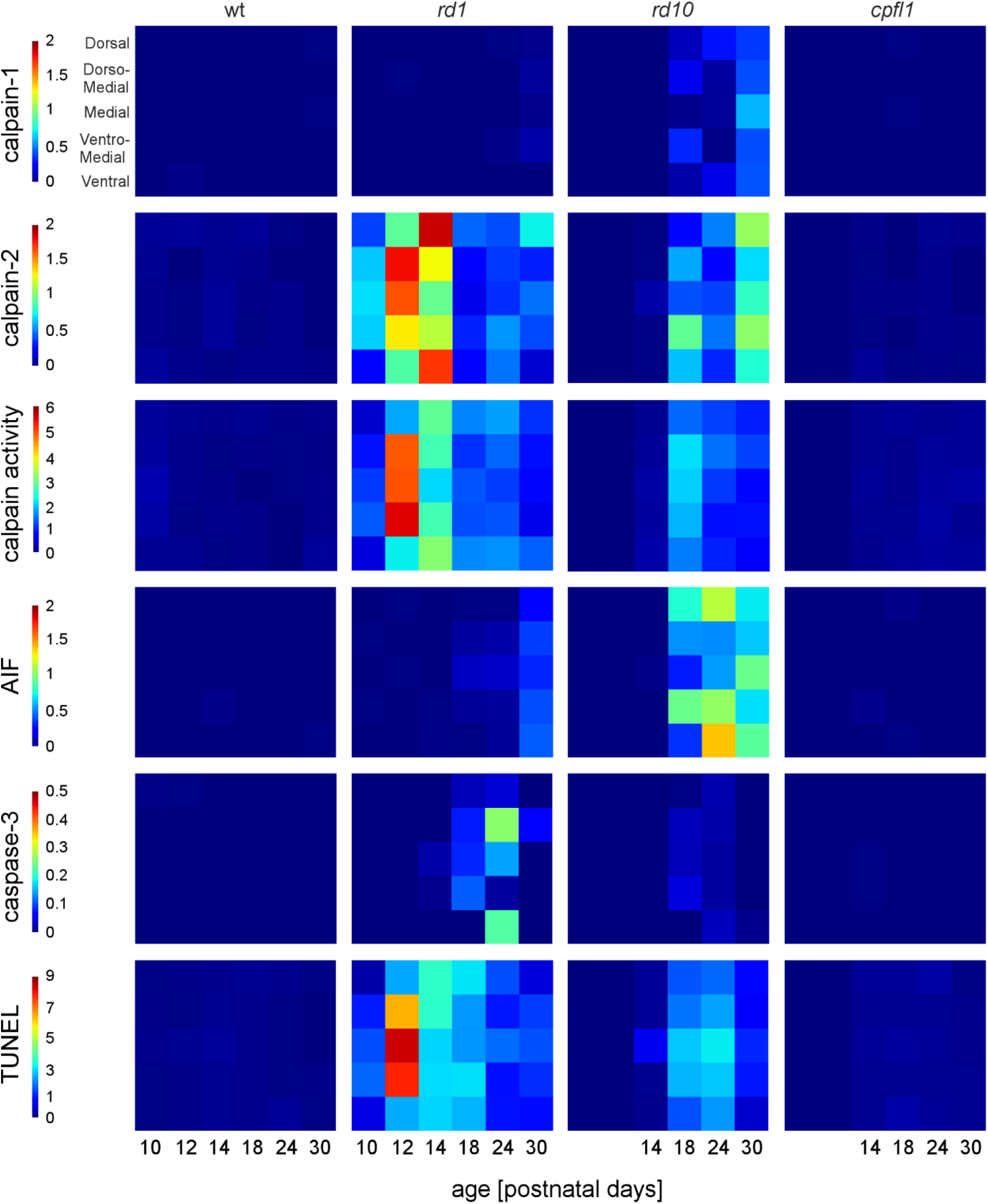
Summary heat maps illustrating the spatio-temporal progression of cell death markers, with image position along the y-axis (dorso-ventral) and time along x-axis (see Methods for details and Figures 11-16 for individual maps). Each element in the heat map is the mean value averaged over the naso-temporal axis. Colours represent number of fluorescent cells in the ONL (per 1,000 μm^2^) for calpain-1, calpain-2, calpain activity, AIF, caspase-3, and TUNEL.

**Figure 11.**
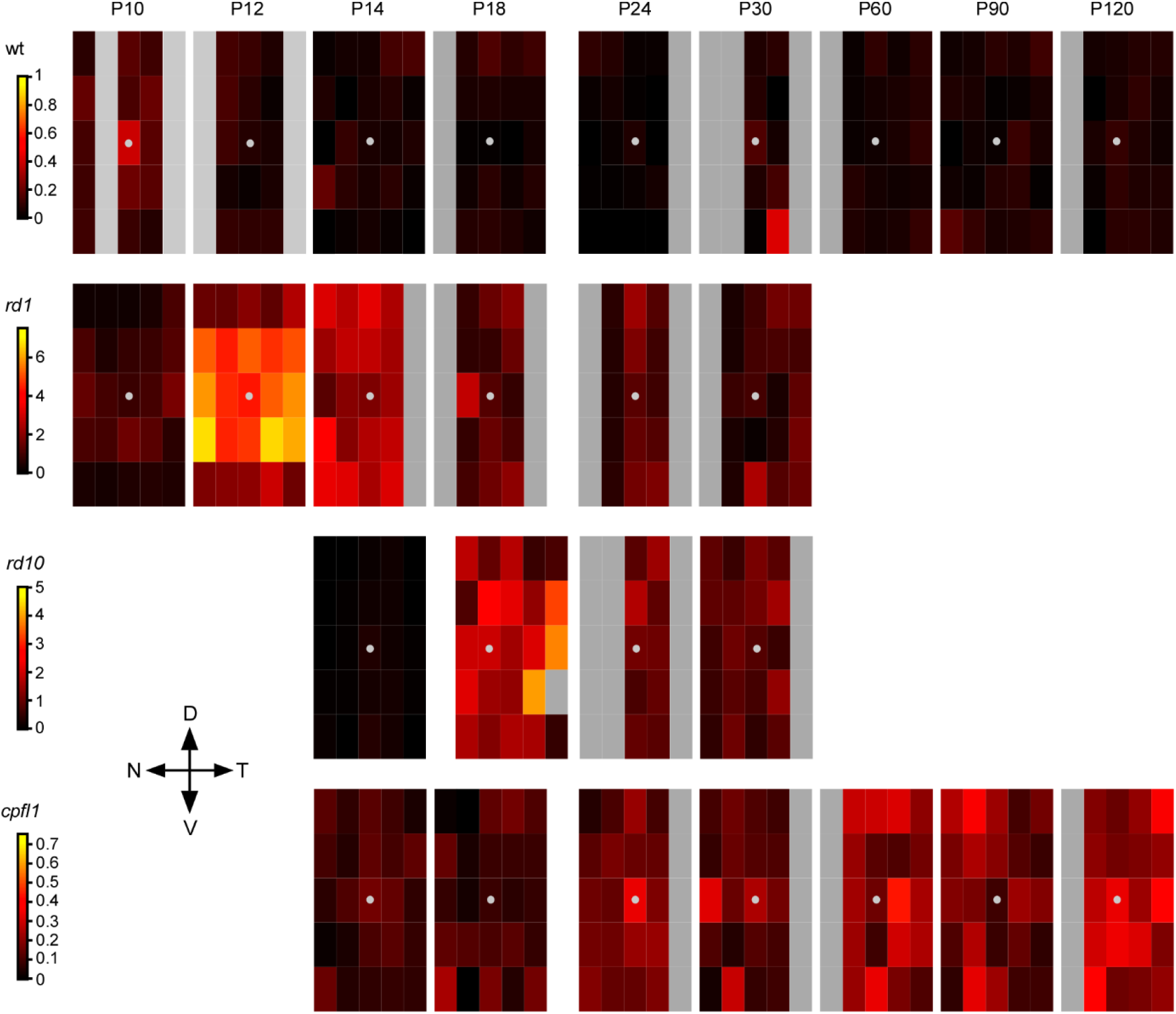
Heat maps for the spatio-temporal progression of calpain activity across the retina and for different mouse lines. Each map represents a time-point, the grey disk the position of the optic nerve head (map origin). Each box in a map measures 1,000 μm by 400 μm, with the colour encoding the average number of fluorescent cells in the ONL (per 1,000 μm^2^). Grey denotes areas that were not examined.

**Figure 12.**
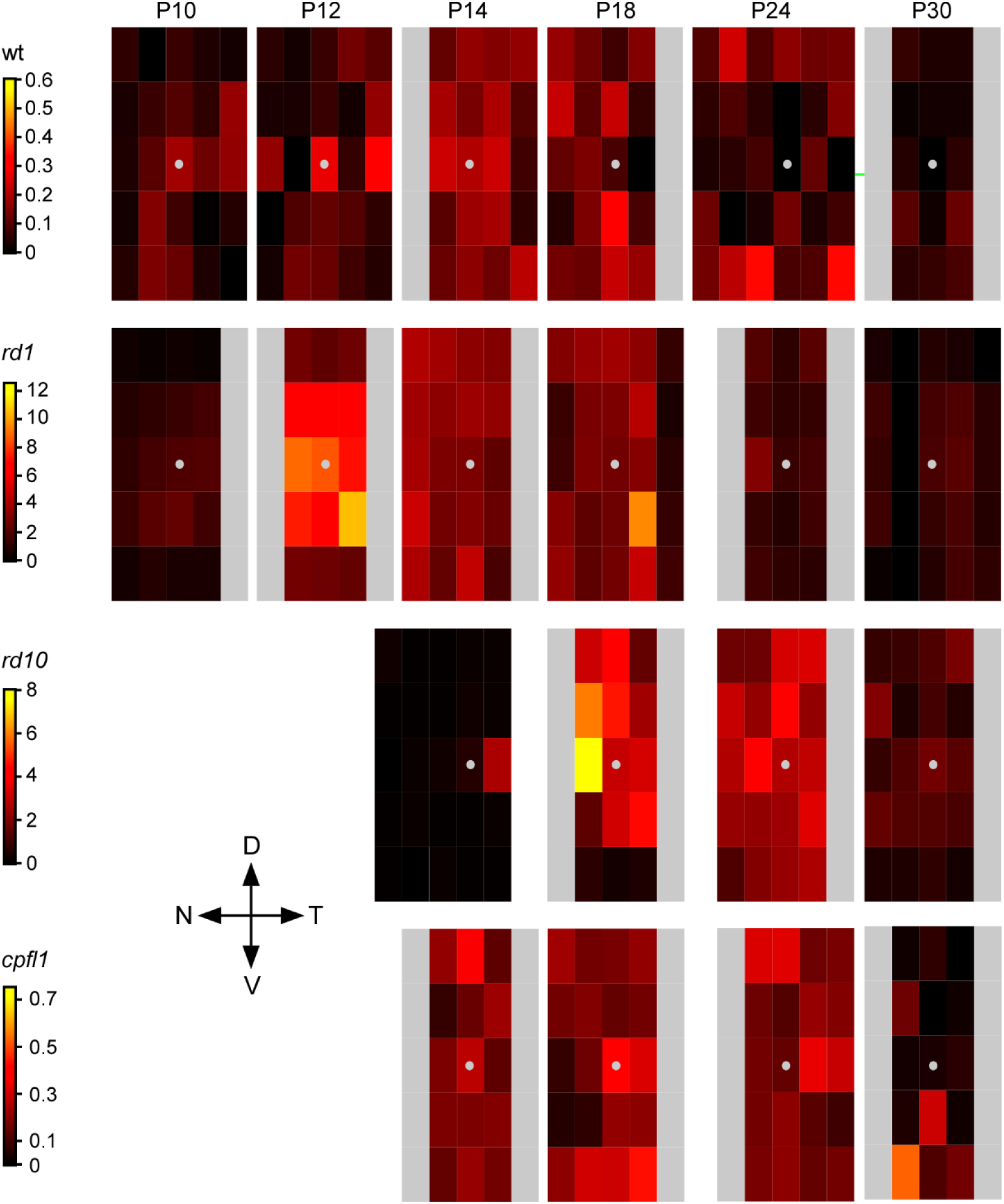
Heat maps for the spatio-temporal progression of cell death (TUNEL) across the retina and for different mouse lines. Each map represents a time-point, the grey disk the position of the optic nerve head (map origin). Each box in a map measures 1,000μm by 400 μm, with the colour encoding the average number of TUNEL positive cells in the ONL (per 1,000 μm^2^). Grey denotes areas that were not examined.

**Figure 13.**
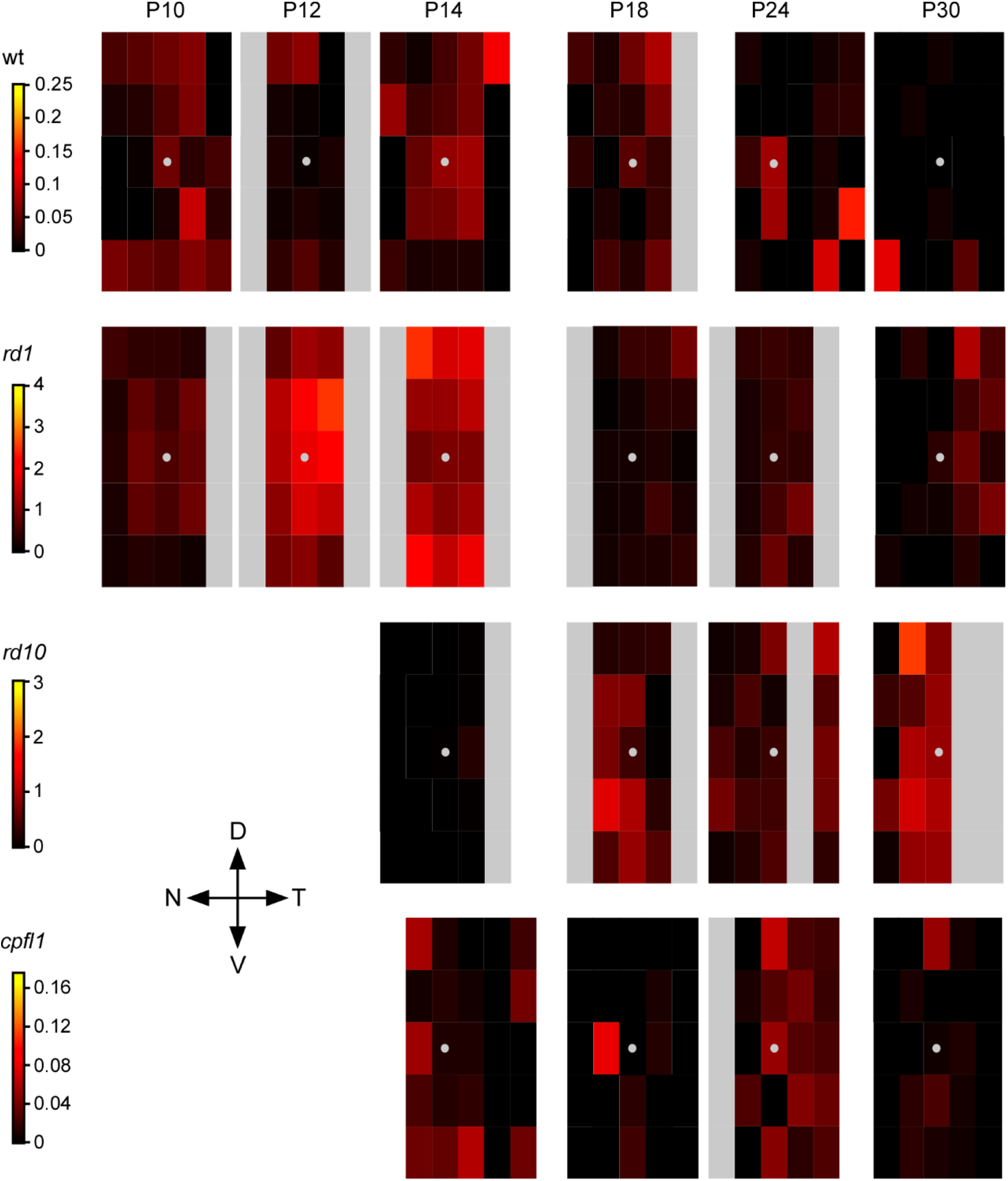
Heat maps for the spatio-temporal progression of calpain-2 activation across the retina and for different mouse lines. Each map represents a time-point, the grey disk the position of the optic nerve head (map origin). Each box in a map measures 1,000 μm by 400 μm, with the colour encoding the average number of fluorescent cells in the ONL (per 1,000 μm^2^). Grey denotes areas that were not examined.

**Figure 14.**
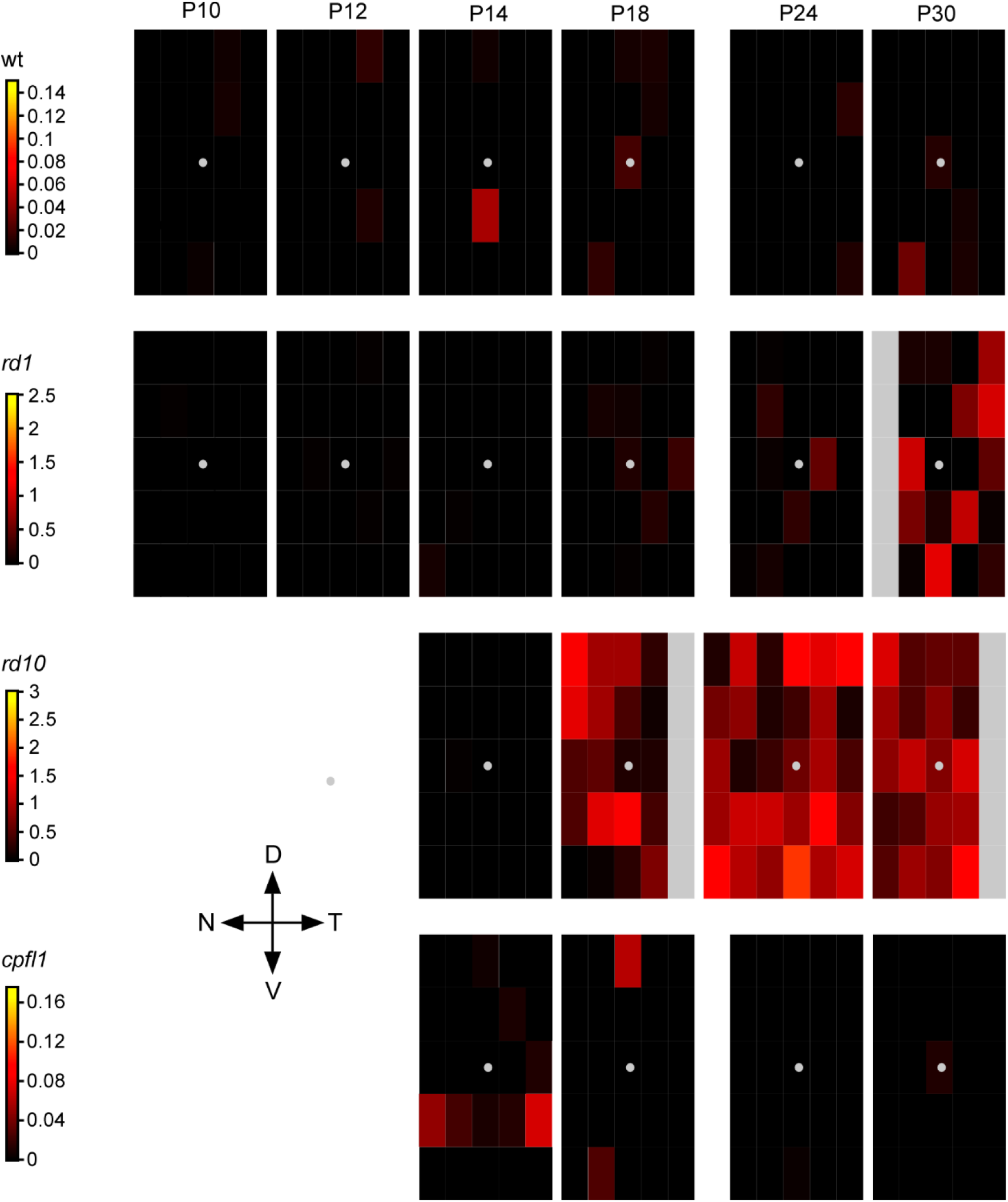
Heat maps for the spatio-temporal progression of AIF immunoreactivity across the retina and for different mouse lines. Each map represents a time-point, the grey disk the position of the optic nerve head (map origin). Each box in a map measures 1,000 μm by 400 μm, with the colour encoding the average number of fluorescent cells in the ONL (per 1,000 μm^2^). Grey denotes areas that were not examined.

**Figure 15.**
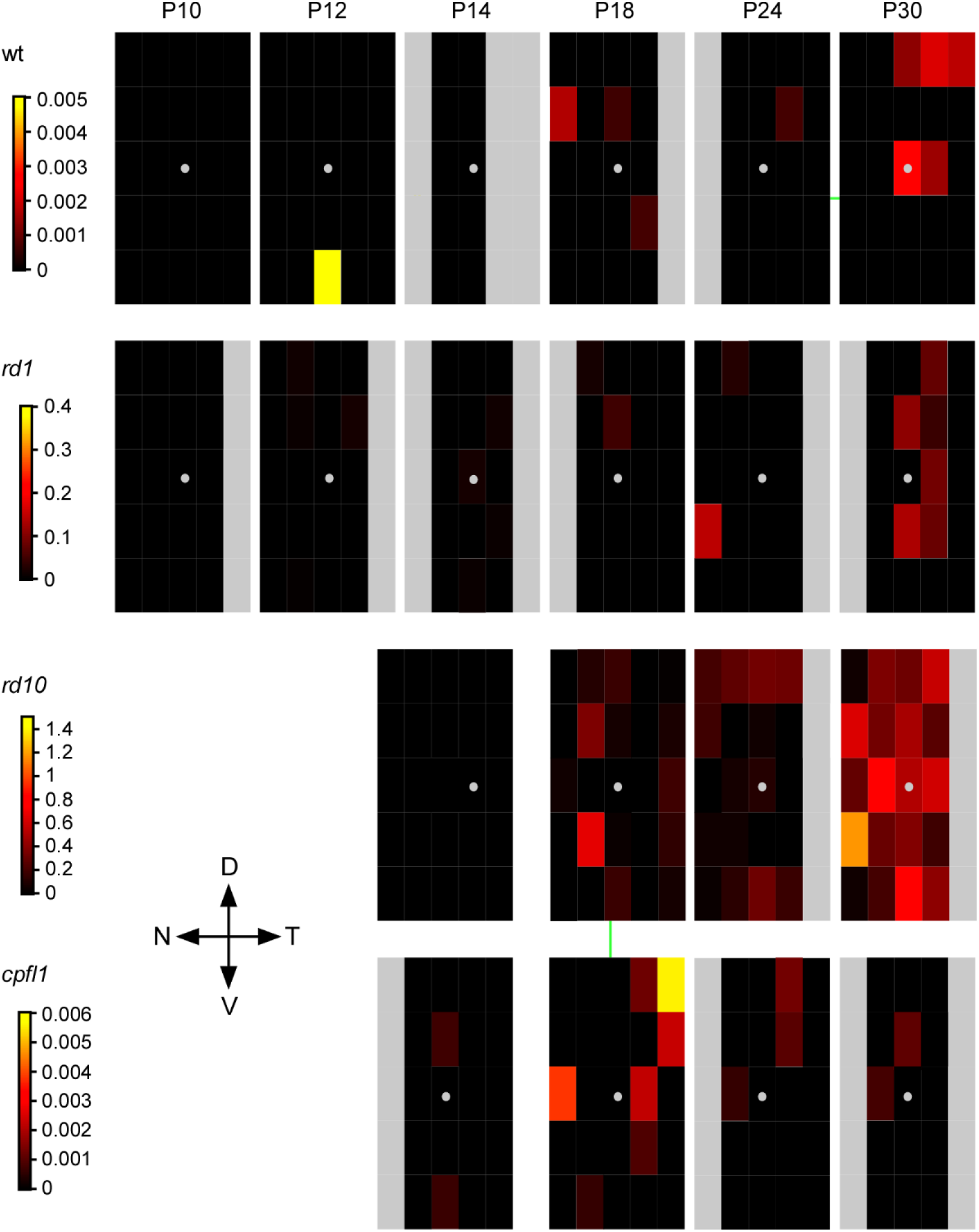
Heat maps for the spatio-temporal progression of calpain-1 activation across the retina and for different mouse lines. Each map represents a time-point, the grey disk the position of the optic nerve head (map origin). Each box in a map measures 1,000 μm by 400 μm, with the colour encoding the average number of fluorescent cells in the ONL (per 1,000 μm^2^). Grey denotes areas that were not examined.

**Figure 16.**
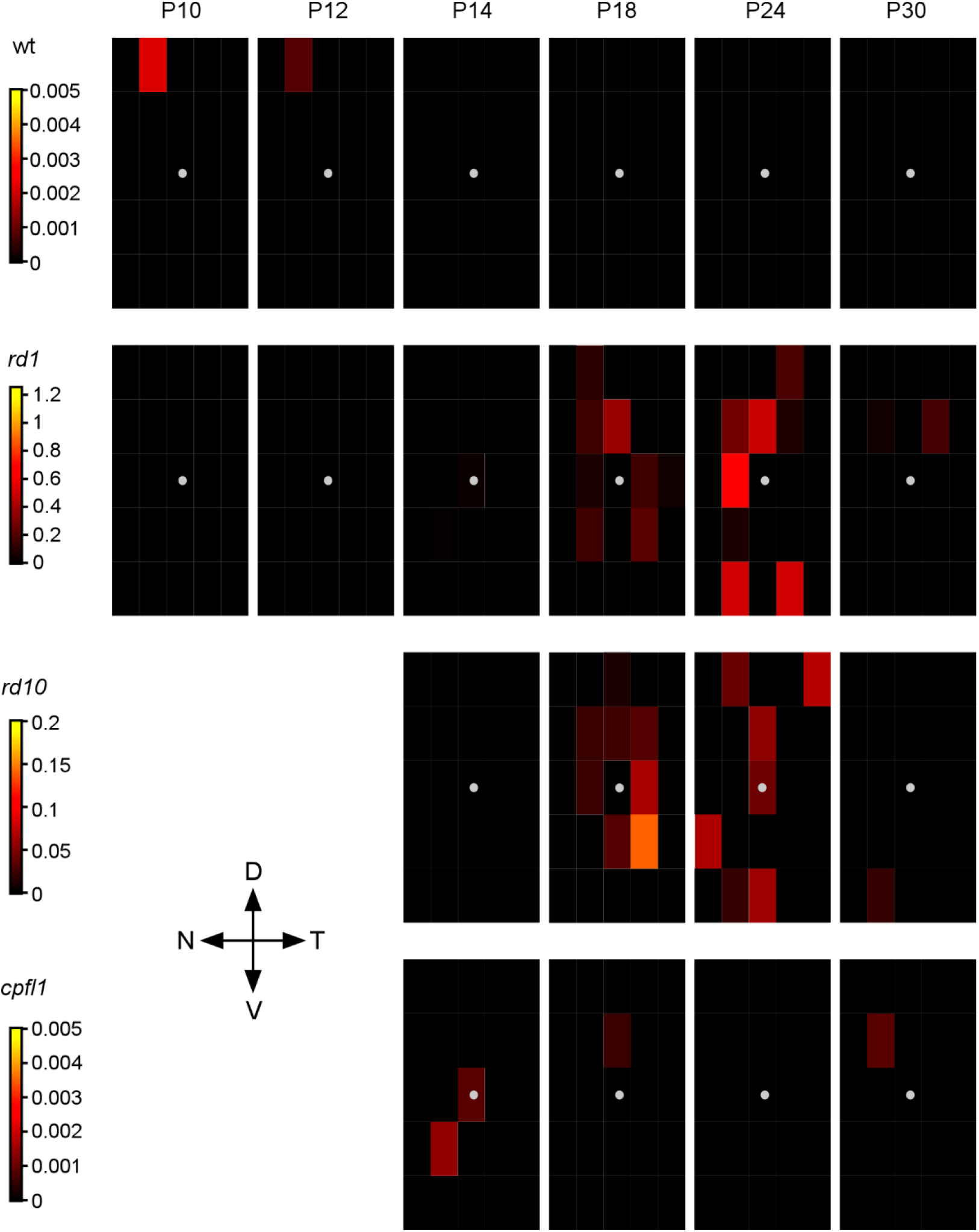
Heat maps for the spatio-temporal progression of caspase-3 activity across the retina and for different mouse lines. Each map represents a time-point, the grey disk the position of the optic nerve head (map origin). Each box in a map measures 1,000 μm by 400 μm, with the colour encoding the average number of fluorescent cells in the ONL (per 1,000 μm^2^). Grey denotes areas that were not examined.

Indeed, the spatial analysis revealed that degeneration did not occur uniformly across the retina of a given time-point: For instance, in *rd1* animals, the peak in calpain activity at P12 first occurred in the central retina and then at P14 in the peripheral retina (Fig. 11, 2^nd^ row). This sequence was also visible in the spatio-temporal maps of *rd1* for calpain activity, TUNEL, and calpain-2 (Fig. 10, 2^nd^ column). A similar trend, though somewhat less clear, was observed in *rd10* retina (Fig. 10, 3^rd^ column; Figs. 11-16, 3^rd^ row). Therefore, this spatial inhomogeneity of cell death markers for a given time-point explains the broad distribution of the data points, at least in part (*cf*. Figs. 3,4,6-8).

### Probabilistic models infer sequences of markers

When comparing the spatio-temporal heat maps for different markers, we noticed that markers within the same retinal region seemed to appear in a certain temporal sequence that differed and/or was delayed between mouse lines. For example, in *rd1* central retina, the peaks of calpain-2, calpain activity, and TUNEL staining appeared to coincide (∼P12), followed by smaller peaks of caspase-3 (P24) and AIF (P30 or later). In *rd10*, the broader “peaks” of calpain-2, calpain activity, and TUNEL also coincided, yet they were more spread out (from P18 to P30) and – as opposed to *rd1* – associated with minor elevations of AIF- and calpain-1, but less so with caspase-3.

To more quantitatively identify the time-points at which each cell death markers peaked in each mouse line, we fitted a set of probabilistic models using Gaussian processes (Fig. 17; for details, see Methods). These models infer the mean and standard deviation of the number of labelled ONL cells over time. To estimate the time at which each marker peaked, we drew 10,000 samples from the posterior of each fitted Gaussian process model and identified the maxima in each sample. The distribution of these maxima indicated the likely peak of each marker (Fig. 18). From these distributions, we identified the most likely sequences of molecular markers in each disease model with a cut-off point of 5% (Fig. 18).

**Figure 17.**
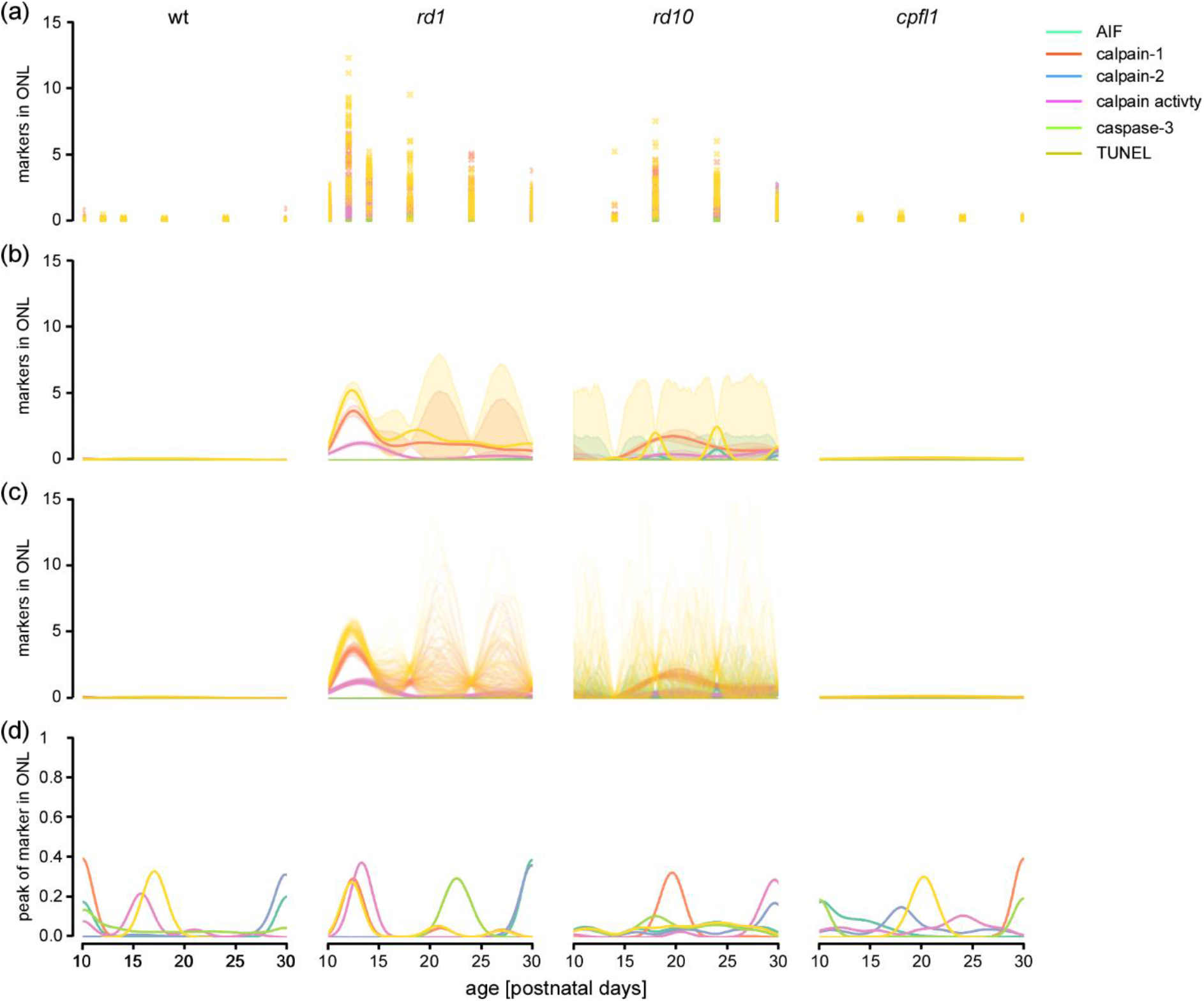
Gaussian process model for the kinetics of cell death related processes. **A**, Parameters associated with photoreceptor cell death as measured in the ONL (per 1,000 μm^2^). **B**, Gaussian Process (GP) models with radial basis function kernel and additive Gaussian noise, fitted to observations to each cross of marker and mouse line. Observations are square root transformed prior to fitting, and after fitting are squared to return the model to its original state. 5% and 95% confidence intervals are estimated using a bootstrap of 1k posterior samples, excluding the additive likelihood noise. **C**, 100 of the 1k posterior samples from the GPs. **D**, Density of the maxima of the GP posterior samples for each marker.

**Figure 18.**
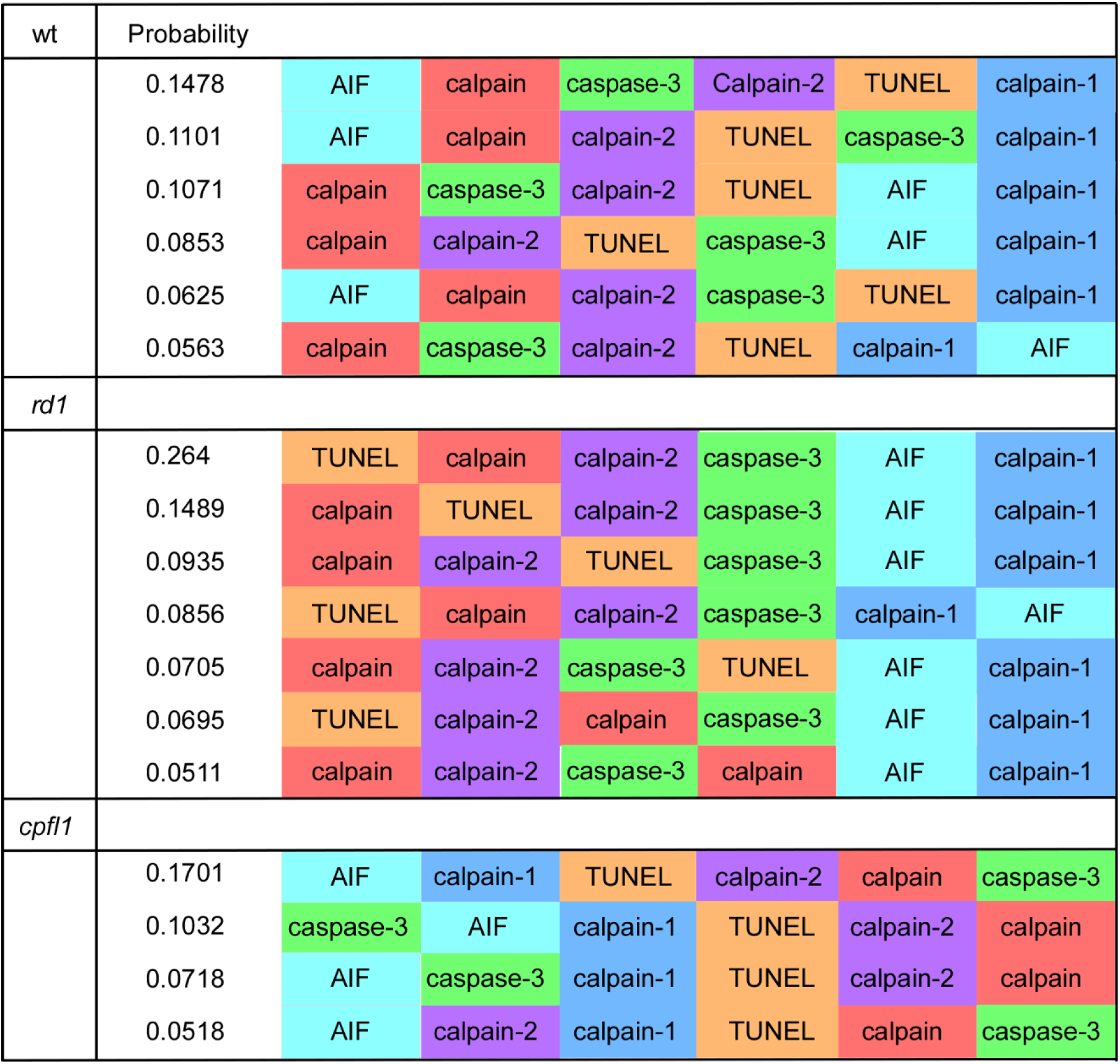
Estimation of marker sequences, using probabilistic Gaussian Process bootstrap. Marker sequences from probabilistic Gaussian Process bootstrap where the probability of observing the sequence was estimated to be greater than 0.05. Probabilities estimated from 1,000 samples for each marker, in each mouse line.

The overall levels of all markers were low for wt and *cpfl1* animals, causing a relatively low level of confidence in the model predictions. In wt mice, calpain activity preceded calpain-2, followed by TUNEL and calpain-1, while in *cpfl1*, AIF was predicted to precede calpain-1 before TUNEL, calpain-2 and calpain activity. In the *rd1* situation there was more confidence in the inferred molecular sequences, showing increased calpain activity, calpain-2-, and TUNEL-positive cells to occur first in the sequence, while AIF and calpain-1 were predicted to always be the last two markers (Fig. 18A). In *rd10*, no sequence had a probability greater than 0.05 (Fig. 19). Taken together, the GP models distinguished between the clear molecular sequences in the *rd1*, compared to the more ambiguous progression in *cpfl1* and *rd10*.

**Figure 19.**
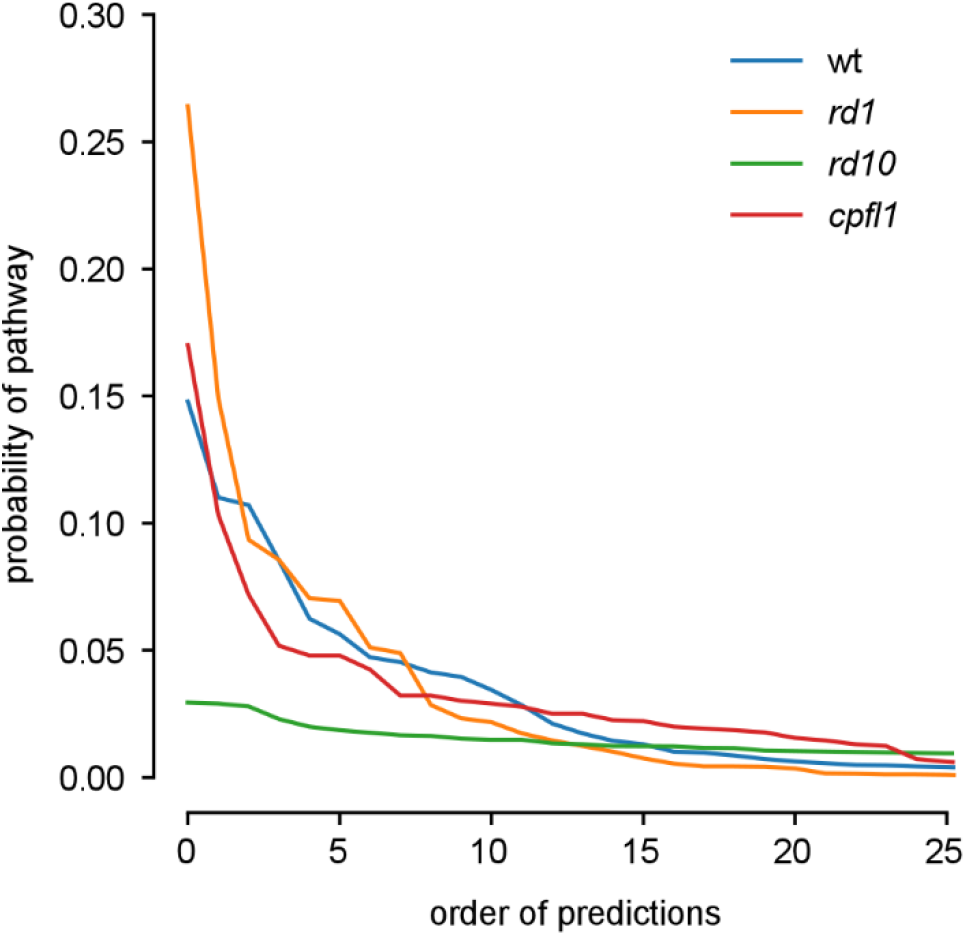
Probability of each marker sequence prediction. Sequence is defined to be the expected order of the peaks for each marker from probabilistic Gaussian Process bootstrap. The probability of each prediction for the order of markers is shown for each mouse line.

## 4. Discussion

Here, we show that activation of calpain-2, but not calpain-1, is strongly correlated with photoreceptor cell death. We show that caspase-3, while not involved in primary rod photoreceptor death, may be responsible for secondary cone degeneration. Our data suggests that peaks for calpain activity, calpain-2, and TUNEL are correlated in space and time starting in the centre of the retina before spreading to the periphery following the onset of degeneration. Modelling results are consistent with the execution of two distinct cell death mechanisms in the *rd1* retina: non-apoptotic cell death during primary rod degeneration and “classical” caspase-driven apoptosis during secondary cone degeneration.

### Ca^2^U dysregulation and cell death

[Ca^2+^] dysregulation occurs in photoreceptors with *Pde6* gene mutations, typically resulting in higher Ca^2^-concentrations (Kulkarni et al., 2016). This rise in [Ca^2^] would lead to increased activity of the plasma membrane NCKX, an exchanger responsible for maintaining appropriate Ca^2^-levels in photoreceptor outer segments (Jensen, Buckby, & Empson, 2004).

NCKX is driven by a K^+^ gradient, which in turn is maintained by Na^+^/K^+^-ATPases. Elsewhere in the cell, low [Ca^2+^] is ensured by specific Ca^2+^-ATPases, which have the highest ATP affinity of all ATPases (Bruce, 2017). This may explain why low [Ca^2+^] can be maintained even when most of the general cellular ATP has become depleted (Bruce, 2017). As necrosis is a passive process that does not require ATP, it has long been suspected that the levels of ATP in the cell are a determining factor for whether, under stress, the cell chooses apoptosis or necrosis to complete cell death (Criddle et al., 2007; Leist et al., 1997; Tsujimoto, 1997). The high levels of Ca^2+^ needed to produce the observed calpain-2 activation could indicate that previously any remaining ATP was already used up by Ca^2+^-ATPases. Hence, a cell death pathway not dependent on ATP (such as necrosis) would seem to be the only pathway left for the photoreceptors to avail of. This concept would agree with the previous finding that the activities of PARP, HDAC, and PKG are also upregulated in models of retinal dystrophies, promoting the execution of a non-apoptotic form of cell death (Arango-Gonzalez et al., 2014; Paquet-Durand et al., 2009; Vighi et al., 2018). Especially activation of PARP has been linked to non-apoptotic cell death, such as parthanatos (David et al.,2009; Galluzzi et al., 2018), while activation of PKG may trigger the execution of anoikis (Hou et al., 2006).

It is tempting to speculate on what could cause such a loss of ATP. A relatively straightforward explanation might be that high cGMP levels caused by *Pde6* mutations, lead to the constitutive over-activation of CNG channels, and thus a constant need for ATP-dependent ion extrusion, which eventually overburdens the photoreceptor capacity to produce ATP. In addition, or alternatively, the down-stream activation of PARP (Paquet-Durand et al., 2007; Sahaboglu et al., 2016) would increase the consumption of NAD^+^ and indirectly decrease ATP-levels (David et al., 2009)

### Secondary cone death driven by apoptosis in rd1

While the majority of *rd1* rods die between P12 and P14, some cells, mostly cones, remain in the outer retina beyond P18. Interestingly, we did find that a substantial cohort of the remaining cones were also positive for caspase-3 in the *rd1* mouse at P24. This would suggest that, while the primary rod degeneration is non-apoptotic, secondary cone degeneration is, in fact, an apoptotic process. In this context, it is also notable that calpain-1 activation was seen in *rd10* retina, but not in *rd1* (or *cpfl1*). Since calpain-1 has been implicated in neuroprotective mechanisms, the remaining photoreceptors may use calpain-1 to prolong their survival (discussed in Baudry & Bi, 2016). When compared to *rd1*, the somewhat slower progression of the *rd10* degeneration may give the cells more time to activate such endogenous calpain-1 dependent protection.

### Spatial occurrences of markers, and progression of cell death

Within the *rd1* mouse retina, a clear centre-to-periphery progression can be seen for calpain activity, calpain-2 up-regulation, and TUNEL staining. This suggests that the activity of calpain (likely calpain-2) drives the advance of cell death across the retina. The centre-to-periphery wave of cell death is well established and thought to follow cell maturation patterns in the *rd1* moose (Noell, 1958). Cell death patterns seen in the *rd10* retina are less well-known but may begin in the mid-periphery and work simultaneously towards and away from the optic nerve (Barhoum et al., 2008; Strettoi & Pignatelli, 2000). In our examination of the *rd10* mouse, a centre to peripheral gradient was indeed seen for calpain activity, calpain-2, and TUNEL, although this gradient was not as distinct as that in the *rd1* mouse. This could be due to time-point selection or to the fact that the degeneration seen in the *rd10* mouse is less synchronous than that in the *rd1* mouse.

In the *cpfl1* model, in agreement with previous studies (Trifunović et al., 2010), calpain activity increased overall, and significantly so at P24. However, no clear pattern of marker up-regulation or activation was detectable, as seen in the individual heat maps (Fig. 11-16). This is probably due to the low number of cones in the mouse retina and the stochastic nature of the cell death seen in all mouse models of retinal dystrophies (Clarke et al., 2000). It is also possible that due to differences in the phototransduction machinery cones are more resilient to Ca^2+^ based cell death mechanisms when compared to rods (Vinberg et al., 2018).

### Different cell death mechanisms in Pde6b mutants?

One of the confounding factors that complicate studies into the causative cell death mechanisms in mouse models is the fact that some of the retinal development and maturation takes place during the 2-3 weeks after birth (Gibson et al., 2013; Young, 1984). When investigating hereditary retinal degeneration in early onset, rapid progression mouse models, this will therefore overlap in time with development and developmental cell death, the latter of which is often thought to be governed by classical apoptotic cell death (Sancho-Pelluz et al., 2008). In wild-type mice developmental photoreceptor cell death occurs in two waves around P15 and P25, likely relating to developmental cell death of rods and cones, respectively (Mervin & Stone, 2002) In this context, the differences seen between *rd1* and *rd10*, with their different onset and progression of retinal degeneration, may help to distinguish developmental from mutation-induced cell death and may thus provide for interesting insights into the causative degeneration mechanisms. While both models show high levels of calpain activity, calpain-2, and TUNEL, at ages corresponding to the degeneration peaks (P12 and P18, respectively), there may be differences in the mode of cell death utilised by the photoreceptors. AIF and calpain-1 activation was prevalent in the ONL of the *rd10* mouse, while the *rd1* retina did not show these markers until relatively late (P24), at which point they may be located predominately to cones. This leads us to ask whether different mutations within the two models cause different pathways of cells death. This question is difficult to address at a cellular level; however, at the tissue level it is obvious that *rd1* and *rd10* mutants differ in the onset and speed of retinal degeneration, with a marked delay and slower progression in *rd10* animals. Interestingly, calpain-1 activity has been implicated in neuroprotective mechanisms (Baudry & Bi, 2016). Hence, the somewhat slower progression of *rd10* degeneration may give individual cells more time to activate endogenous calpain-1 dependent protection (as discussed above).

The *rd10* retina also displayed a more marked up-regulation of AIF compared to *rd1*. Since AIF is a mitochondrial protein, and photoreceptor mitochondria in the inner segments grow and mature with post-natal age, it is conceivable that this apparent increase in AIF is entirely due to the later onset of *rd10* degeneration. If correct, then AIF may also be associated with *rd1* degeneration, as has been suggested before (Sanges et al., 2006), but would be more difficult to detect because at the onset of *rd1* degeneration its expression was much lower.

Taken together, the cell death mechanisms seen in primary rod degeneration, in both *rd1* and *rd10* retina, are clearly connected to calpain-2 activation, independent of caspase-3, and likely involve AIF release from mitochondria – hence, indicative of non-apoptotic cell death pathways. Conversely, calpain-1 activation may play a protective role, however, a role which ultimately is not strong enough to save photoreceptors from mutation-induced degeneration.

### Future directions

Here, we adopted a “standard” experimental strategy which emphasises the use of large numbers of samples at a small number of predetermined time-points (Arango-Gonzalez et al., 2014; Kulkarni et al., 2016). Although this approach was useful for inferring the state of the system at particular disease stages, the rigidity of the sampling came at the cost of lower temporal resolution. Where the temporal precision is critical, it would be preferable to adopt a proactive strategy, using GP models to pre-select the optimal time-points for particular hypotheses (Chaloner & Verdinelli, 1995; Lindley, 1956; Pillow, 2016). This should make it possible to infer more accurately the non-linear progression of each marker over time, with fewer samples than would be needed in a predetermined or purely random approach.

In this study, we employed a GP model to identify more precisely the sequence of degenerative events occurring within each cell. This model suited our investigation well as it was able to infer a probabilistic representation of the progression of each marker, and could be fitted to unevenly spaced samples. The principled representation of uncertainty also reflected the noisiness and heterogeneity of the molecular progression. Studies where low levels of total cell death are observed may benefit from using Poisson likelihood models, rather than the Gaussian distributions in our approach (Adams et al., 2009). Implementations of Poisson processes are available in the Gaussian Process library GPy (GPy, 2014). Additionally, these models may not be directly applicable to diseases where cell death is induced by an abrupt, cataclysmic and isolated event. This is because the models that we have presented assume that the statistical properties of the molecular sequences are stationary over time, but there are extensions to GPs which can handle non-stationarity (Snelson et al., 2004).

## Conclusions

Temporal-spatial mapping of degenerative markers in the retina showed that Ca^2+^-dependent calpain activity in hereditary retinal degeneration rises in congress with the degeneration of the tissue. The observed calpain activity was strongly correlated and likely caused by activation of calpain-2, suggesting calpain-2 as an important driver of photoreceptor cell death. We provide further strong evidence for primary rod degeneration being a non-apoptotic mechanism, even though, at least in *rd1* retina, the delayed appearance of caspase-3 activation in cones suggests apoptosis as a driver of secondary cone degeneration.

These results demonstrate the complexity of cell death mechanisms in hereditary retinal degeneration and highlight the importance of classifying the relevant cell death mechanisms, in the various stages of disease progression. Notably, the confirmation and identification of Ca^2+^-dysregulation and calpain-2 as components of a pathway driving photoreceptor degeneration may guide the development of future therapeutics.

## Acknowledgements

We thank Sylvia Bolz, Norman Rieger, and Gordon Eske for excellent technical support and guidance. We are grateful to Bernd Wissinger for providing us with the *rd8* genotyping protocol. We also thank Philipp Hennig for commentary and feedback on the GP modelling.

## List of abbreviations

Ca^2+^: calcium
AIF: apoptosis inducing factor
RD: retinal dystrophies
RP: retinitis pigmentosa
Pde6: phosphodiesterase 6
cGMP: cyclic guanosine monophosphate
Na^+^: sodium
P: postnatal
DTT: 1,4-Dithiothreitol
CRB: calpain reaction buffer
PBS: phosphor buffered saline
TUNEL: Terminal deoxynucleotidyl transferase dUTP nick end labeling
GPs: Gaussian Processes
ONL: outer nuclear layer
NCKX: Na+/Ca2+-K+ exchangers

## Declarations

### Ethics approval and consent to participate

All procedures were performed in accordance with the law on animal protection issued by the German Federal Government (Tierschutzgesetz) and approved by the institutional animal welfare office of the University of Tübingen.

### Consent for publication

N/A

### Availability of data and material

The dataset(s) supporting the conclusions of this article is(are) available in the Zenodo repository, 10.5281/zenodo.2571443 and https://zenodo.org/record/2571443#.XGqNmehKjcs. The manuscript is available in pre-print form at https://doi.org/10.1101/554733

## Authors contributions

MP, TE and FPD designed the experiments which were carried out by MP. Initial analysis was carried out by MP with LR contributing further analysis and the Gaussian process modelling and helped initially format the spreadsheets. PB helped with statistical analysis, TS helped scrutinise the data and create the figures. All authors reviewed and helped writing the manuscript.

## References

Adams, R. P., Murray, I., & MacKay, D. J. C. (2009). Tractable nonparametric Bayesian inference in Poisson processes with Gaussian process intensities. In Proceedings of the 26th Annual International Conference on Machine Learning (pp. 9–16). ACM.

Arango-Gonzalez, B., Trifunović, D., Sahaboglu, A., Kranz, K., Michalakis, S., Farinelli, P., … Janssen-Bienhold, U. (2014). Identification of a common non-apoptotic cell death mechanism in hereditary retinal degeneration. PloS One, 9(11), e112142.

Bano, D., & Prehn, J. H. M. (2018). Apoptosis-Inducing Factor (AIF) in Physiology and Disease: The Tale of a Repented Natural Born Killer. EBioMedicine.

Barhoum, R., Martinez-Navarrete, G., Corrochano, S., Germain, F., Fernández-Sánchez, L., De la Rosa, E. J., … Cuenca, N. (2008). Functional and structural modifications during retinal degeneration in the rd10 mouse. Neuroscience, 155(3), 698–713.

Baudry, M., & Bi, X. (2016). Calpain-1 and calpain-2: the yin and yang of synaptic plasticity and neurodegeneration. Trends in Neurosciences, 39(4), 235–245.

Behrens, C., Schubert, T., Haverkamp, S., Euler, T., & Berens, P. (2016). Connectivity map of bipolar cells and photoreceptors in the mouse retina. Elife, 5, e20041.

Bertelsen, M., Jensen, H., Bregnhøj, J. F., & Rosenberg, T. (2014). Prevalence of generalized retinal dystrophy in Denmark. Ophthalmic Epidemiology, 21(4), 217–223.

Bruce, J. I. E. (2017). Metabolic regulation of the PMCA: role in cell death and survival. Cell Calcium.

Chaloner, K., & Verdinelli, I. (1995). Bayesian experimental design: A review. Statistical Science, 273–304.

Chang, B, Hawes, N. L., Hurd, R. E., Davisson, M. T., Nusinowitz, S., & Heckenlively, J. R. (2002). Retinal degeneration mutants in the mouse. Vision Research, 42(4), 517–525.

Chang, Bo, Grau, T., Dangel, S., Hurd, R., Jurklies, B., Sener, E. C., … Bolz, S. (2009). A homologous genetic basis of the murine cpfl1 mutant and human achromatopsia linked to mutations in the PDE6C gene. Proceedings of the National Academy of Sciences, 106(46), 19581–19586.

Chen, L. Y., Rex, C. S., Casale, M. S., Gall, C. M., & Lynch, G. (2007). Changes in synaptic morphology accompany actin signaling during LTP. Journal of Neuroscience, 27(20), 5363–5372.

Clarke, G., Collins, R. A., Leavitt, B. R., Andrews, D. F., Hayden, M. R., Lumsden, C. J., & McInnes, R. R. (2000). A one-hit model of cell death in inherited neuronal degenerations. Nature, 406(6792), 195.

Criddle, D. N., Gerasimenko, J. V., Baumgartner, H. K., Jaffar, M., Voronina, S., Sutton, R., … Gerasimenko, O. V. (2007). Calcium signalling and pancreatic cell death: apoptosis or necrosis? Cell Death and Differentiation, 14(7), 1285.

Croall, D. E., & Ersfeld, K. (2007). The calpains: modular designs and functional diversity. Genome Biology, 8(6), 218.

Danciger, M., Blaney, J., Gao, Y. Q., Zhao, D. Y., Heckenlively, J. R., Jacobson, S. G., & Farber, D. B. (1995). Mutations in the PDE6B Gene in Autosomal Recessive Retinitis Pigmentosa. Genomics, 30(1), 1–7. https://doi.org/10.1006/geno.1995.0001

David, K. K., Andrabi, S. A., Dawson, T. M., & Dawson, V. L. (2009). Parthanatos, a messenger of death. Frontiers in Bioscience (Landmark Edition), 14, 1116.

Doonan, F., Donovan, M., & Cotter, T. G. (2005). Activation of multiple pathways during photoreceptor apoptosis in the rd mouse. Investigative Ophthalmology & Visual Science, 46(10), 3530–3538.

Galluzzi, L., Vitale, I., Aaronson, S. A., Abrams, J. M., Adam, D., Agostinis, P., … Kroemer, G. (2018). Molecular mechanisms of cell death: recommendations of the Nomenclature Committee on Cell Death 2018. Cell Death & Differentiation, 25(3), 486–541. https://doi.org/10.1038/s41418-017-0012-4

Gibson, R., Fletcher, E. L., Vingrys, A. J., Zhu, Y., Vessey, K. A., & Kalloniatis, M. (2013). Functional and neurochemical development in the normal and degenerating mouse retina. Journal of Comparative Neurology, 521(6), 1251–1267.

Goll, D E. (1995). Properties and biological regulation of the calpain system. Expression of Tissue Proteinases and Regulation of Protein Degradation as Related to Meat Quality, 47–68.

Goll, Darrel E, Thompson, V. F., Li, H., Wei, W. E. I., & Cong, J. (2003). The calpain system. Physiological Reviews, 83(3), 731–801.

Goñi-Oliver, P., Lucas, J. J., Avila, J., & Hernández, F. (2007). N-terminal Cleavage of GSK-3 by Calpain a new form of GSK-3 regulation. Journal of Biological Chemistry, 282(31), 22406–22413.

GPy. (2014). GPy: A gaussian process framework in python. Retrieved from http://github.com/SheffieldML/GPy

Hamel, C. (2006). Retinitis pigmentosa. Orphanet Journal of Rare Diseases, 1(1), 40.

Hartong, D. T., Berson, E. L., & Dryja, T. P. (2006). Retinitis pigmentosa. The Lancet, 368(9549), 1795–1809.

Hou, Y., Wong, E., Martin, J., Schoenlein, P. V, Dostmann, W. R., & Browning, D. D. (2006). A role for cyclic-GMP dependent protein kinase in anoikis. Cellular Signalling, 18(6), 882–888.

Jensen, T. P., Buckby, L. E., & Empson, R. M. (2004). Expression of plasma membrane Ca2+ ATPase family members and associated synaptic proteins in acute and cultured organotypic hippocampal slices from rat. Developmental Brain Research, 152(2), 129– 136.

Jeon, C.-J., Strettoi, E., & Masland, R. H. (1998). The major cell populations of the mouse retina. Journal of Neuroscience, 18(21), 8936–8946.

Keeler, C. (1966). Retinal degeneration in the mouse is rodless retina. Journal of Heredity, 57(2), 47–50.

Khorchid, A., & Ikura, M. (2002). How calpain is activated by calcium. Nature Structural and Molecular Biology, 9(4), 239.

Kohl, S., Coppieters, F., Meire, F., Schaich, S., Roosing, S., Brennenstuhl, C., … Lukowski, R. (2012). A nonsense mutation in PDE6H causes autosomal-recessive incomplete achromatopsia. The American Journal of Human Genetics, 91(3), 527–532.

Kraupp, B. G., Ruttkay Nedecky, B., Koudelka, H., Bukowska, K., Bursch, W., & Schulte Hermann, R. (1995). In situ detection of fragmented DNA (TUNEL assay) fails to discriminate among apoptosis, necrosis, and autolytic cell death: a cautionary note. Hepatology, 21(5), 1465–1468.

Krizaj, D., & Copenhagen, D. R. (2002). Calcium regulation in photoreceptors. Frontiers in Bioscience: A Journal and Virtual Library, 7, d2023.

Kulkarni, M., Trifunović, D., Schubert, T., Euler, T., & Paquet-Durand, F. (2016). Calcium dynamics change in degenerating cone photoreceptors. Human Molecular Genetics, 25(17), 3729–3740.

Leist, M., Single, B., Castoldi, A. F., Kühnle, S., & Nicotera, P. (1997). Intracellular adenosine triphosphate (ATP) concentration: a switch in the decision between apoptosis and necrosis. Journal of Experimental Medicine, 185(8), 1481–1486.

Lindley, D. V. (1956). On a Measure of the Information Provided by an Experiment. The Annals of Mathematical Statistics, 27(4), 986–1005. Retrieved from http://www.jstor.org/stable/2237191

Liu, X., Van Vleet, T., & Schnellmann, R. G. (2004). The role of calpain in oncotic cell death. Annu. Rev. Pharmacol. Toxicol., 44, 349–370.

Mank, M., Reiff, D. F., Heim, N., Friedrich, M. W., Borst, A., & Griesbeck, O. (2006). A FRET-based calcium biosensor with fast signal kinetics and high fluorescence change. Biophysical Journal, 90(5), 1790–1796.

Mattapallil, M. J., Wawrousek, E. F., Chan, C.-C., Zhao, H., Roychoudhury, J., Ferguson, T. A., & Caspi, R. R. (2012). The Rd8 mutation of the Crb1 gene is present in vendor lines of C57BL/6N mice and embryonic stem cells, and confounds ocular induced mutant phenotypes. Investigative Ophthalmology & Visual Science, 53(6), 2921–2927.

Mazumder, S., Plesca, D., & Almasan, A. (2008). Caspase-3 activation is a critical determinant of genotoxic stress-induced apoptosis. In Apoptosis and Cancer (pp. 13–21). Springer.

McLaughlin, M. E., Sandberg, M. A., Berson, E. L., & Dryja, T. P. (1993). Recessive mutations in the gene encoding the β–subunit of rod phosphodiesterase in patients with retinitis pigmentosa. Nature Genetics, 4(2), 130.

Mervin, K., & Stone, J. (2002). Developmental death of photoreceptors in the C57BL/6JMouse: association with retinal function and self-protection. Experimental Eye Research, 75(6), 703–713.

Michalakis, S., Becirovic, E., & Biel, M. (2018). Retinal cyclic nucleotide-gated channels: from pathophysiology to therapy. International Journal of Molecular Sciences, 19(3), 749.

Nemova, N. N., Lysenko, L. A., & Kantserova, N. P. (2010). Proteases of the calpain family: Structure and functions. Russian Journal of Developmental Biology, 41(5), 318–325. https://doi.org/10.1134/S1062360410050073

Noell, W. K. (1958). Studies on visual cell viability and differentiation. Annals of the New York Academy of Sciences, 74(1), 337–361.

Orrenius, S., Zhihotovsky, B., & Nicotera, P. (2003). Regulation of cell death: the calcium-apoptosis link. Nature Rev Cell Biol 2003; 4: 552–65.

Paquet-Durand, Francois, Azadi, S., Hauck, S. M., Ueffing, M., van Veen, T., & Ekström, P. (2006). Calpain is activated in degenerating photoreceptors in the rd1 mouse. Journal of Neurochemistry, 96(3), 802–814.

Paquet-Durand, François, Hauck, S. M., Van Veen, T., Ueffing, M., & Ekström, P. (2009). PKG activity causes photoreceptor cell death in two retinitis pigmentosa models. Journal of Neurochemistry, 108(3), 796–810.

Paquet-Durand, François, Johnson, L., & Ekström, P. (2007). Calpain activity in retinal degeneration. Journal of Neuroscience Research, 85(4), 693–702.

Pillow, J.W., P. M. (2016). Adaptive Bayesian methods for closed-loop neurophysiology. In Closed Loop Neuroscience.

Remmer, M. H., Rastogi, N., Ranka, M. P., & Ceisler, E. J. (2015). Achromatopsia: a review. Current Opinion in Ophthalmology, 26(5).

Rizzuto, R., Pinton, P., Ferrari, D., Chami, M., Szabadkai, G., Magalhaes, P. J., … Pozzan, T. (2003). Calcium and apoptosis: facts and hypotheses. Oncogene, 22(53), 8619.

Sahaboglu, A., Barth, M., Secer, E., Del Amo, E. M., Urtti, A., Arsenijevic, Y., … Paquet-Durand, F. (2016). Olaparib significantly delays photoreceptor loss in a model for hereditary retinal degeneration. Scientific Reports, 6, 39537.

Sancho-Pelluz, J., Arango-Gonzalez, B., Kustermann, S., Romero, F. J., van Veen, T., Zrenner, E., … Paquet-Durand, F. (2008). Photoreceptor cell death mechanisms in inherited retinal degeneration. Molecular Neurobiology, 38(3), 253–269.

Sanges, D., Comitato, A., Tammaro, R., & Marigo, V. (2006). Apoptosis in retinal degeneration involves cross-talk between apoptosis-inducing factor (AIF) and caspase-12 and is blocked by calpain inhibitors. Proceedings of the National Academy of Sciences, 103(46), 17366–17371.

Shang, L., Huang, J.-F., Ding, W., Chen, S., Xue, L.-X., Ma, R.-F., & Xiong, K. (2014). Calpain: a molecule to induce AIF-mediated necroptosis in RGC-5 following elevated hydrostatic pressure. BMC Neuroscience, 15(1), 63.

Snelson, E., Ghahramani, Z., & Rasmussen, C. E. (2004). Warped gaussian processes. In Advances in neural information processing systems (pp. 337–344).

Strettoi, E., & Pignatelli, V. (2000). Modifications of retinal neurons in a mouse model of retinitis pigmentosa. Proceedings of the National Academy of Sciences, 97(20), 11020– 11025.

Sundaram, V., Wilde, C., Aboshiha, J., Cowing, J., Han, C., Langlo, C. S., … Bainbridge, J. W. (2014). Retinal structure and function in achromatopsia: implications for gene therapy. Ophthalmology, 121(1), 234–245.

Suzuki, K., Hata, S., Kawabata, Y., & Sorimachi, H. (2004). Structure, activation, and biology of calpain. Diabetes, 53(suppl 1), S12–S18.

Thiadens, A. A. H. J., den Hollander, A. I., Roosing, S., Nabuurs, S. B., Zekveld-Vroon, R. C., Collin, R. W. J., … Strom, T. M. (2009). Homozygosity mapping reveals PDE6C mutations in patients with early-onset cone photoreceptor disorders. The American Journal of Human Genetics, 85(2), 240–247.

Trifunović, D., Dengler, K., Michalakis, S., Zrenner, E., Wissinger, B., & Paquet-Durand, F. (2010). CGMP-dependent cone photoreceptor degeneration in the cpfl1 mouse retina. Journal of Comparative Neurology, 518(17), 3604–3617. https://doi.org/10.1002/cne.22416

Tsujimoto, Y. (1997). Apoptosis and necrosis: intracellular ATP level as a determinant for cell death modes. Cell Death and Differentiation, 4(6), 429.

Vighi, E., Trifunović, D., Veiga-Crespo, P., Rentsch, A., Hoffmann, D., Sahaboglu, A., … Van Den Heuvel, A. (2018). Combination of cGMP analogue and drug delivery system provides functional protection in hereditary retinal degeneration. Proceedings of the National Academy of Sciences, 115(13), E2997–E3006.

Vinberg, F., Chen, J., & Kefalov, V. J. (2018). Regulation of calcium homeostasis in the outer segments of rod and cone photoreceptors. Progress in Retinal and Eye Research, 67, 87– 101.

Wang, S., Liao, L., Wang, M., Zhou, H., Huang, Y., Wang, Z., … Wang, Y. (2018). Pin1 promotes regulated necrosis induced by glutamate in rat retinal neurons via CAST/Calpain2 pathway. Frontiers in Cellular Neuroscience, 11, 425.

Wei, T., Schubert, T., Paquet-Durand, F., Tanimoto, N., Chang, L., Koeppen, K., … Euler, T. (2012). Light-driven calcium signals in mouse cone photoreceptors. Journal of Neuroscience, 32(20), 6981–6994.

Williams, C. K., & Rasmussen, C. E. (2006). Gaussian processes for machine learning. The MIT Press, 2(3), 4.

Young, R. W. (1984). Cell death during differentiation of the retina in the mouse. Journal of Comparative Neurology, 229(3), 362–373.

